# Multi-modal comparison of molecular programs driving nurse cell death and clearance in *Drosophila melanogaster* oogenesis

**DOI:** 10.1101/2024.03.12.584558

**Authors:** Shruthi Bandyadka, Diane PV Lebo, Albert Mondragon, Sandy B Serizier, Julian Kwan, Jeanne S Peterson, Alexandra Y Chasse, Victoria Jenkins, Anoush Calikyan, Anthony Ortega, Joshua D Campbell, Andrew Emili, Kimberly McCall

## Abstract

The death and clearance of nurse cells is a consequential milestone in *Drosophila melanogaster* oogenesis. In preparation for oviposition, the germline-derived nurse cells bequeath to the developing oocyte all their cytoplasmic contents and undergo programmed cell death. The death of the nurse cells is controlled non-autonomously and is precipitated by epithelial follicle cells of somatic origin acquiring a squamous morphology and acidifying the nurse cells externally. Alternatively, stressors such as starvation can induce the death of nurse cells earlier in mid-oogenesis, manifesting apoptosis signatures, followed by their engulfment by epithelial follicle cells. To identify and contrast the molecular pathways underlying these morphologically and genetically distinct cell death paradigms, both mediated by follicle cells, we compared their genome-wide transcriptional, translational, and secretion profiles before and after differentiating to acquire a phagocytic capability, as well as during well-fed and nutrient-deprived conditions. By coupling the GAL4-UAS system to Translating Ribosome Affinity Purification (TRAP-seq) and proximity labeling (HRP-KDEL) followed by Liquid Chromatography tandem mass-spectrometry, we performed high-throughput screens to identify pathways selectively activated or repressed by follicle cells to employ nurse cell-clearance routines contextually and preferentially. We also integrated two publicly available single-cell RNAseq atlases of the *Drosophila* ovary to define the transcriptomic profiles of follicle cells. In this report, we describe the genes and major pathways identified in the screens and the striking consequences to *Drosophila melanogaster* oogenesis caused by RNAi perturbation of prioritized candidates. To the best of our knowledge, our study is the first of its kind to comprehensively characterize two distinct apoptotic and non-apoptotic cell death paradigms in the same multi-cellular system. Beyond molecular differences in cell death, our investigation may also provide insights into how key systemic trade-offs are made between survival and reproduction when faced with physiological stress.

## Introduction

Oogenesis is a tightly regulated interplay of germline and somatic cells conspiring to produce a viable egg. Like industrial production lines, organisms have evolved to reinforce egg production by adapting quality control measures, such as inspection checkpoints, corrective action, and elimination of defective egg chambers to optimize the health of eggs. Cell death is a requisite means to cull both sub-optimal products of oogenesis, as well as superfluous peripheral tissues that have completed their respective roles in supporting the oocyte on its way to maturation [1].

In *Drosophila melanogaster*, the egg chamber undergoes 14 well-defined stages of development, with significant physical and physiological interactions between the oocyte, the germline-derived Nurse Cells (NCs) and the somatic Follicle Cells (FCs) that form a monolayer encapsulating the germline [2,3]. Multiple regulated cell death events mark the checkpoints during which both the egg’s competence and the suitability of the environment is surveyed. One such crucial checkpoint is encountered in mid-oogenesis during stages 7 through 9, when the decision to invest energy into yolk, chorion, and vitelline membrane deposition is made. As previously reported [4,5], seemingly innocuous physiological stressors such as short-term protein deprivation can induce a 60-fold difference in the rate of oogenesis. This diminished oviposition results in part from the degeneration of egg chambers in mid-oogenesis, which can be readily identified by the condensed and fragmented appearance of NC nuclei and the enlargement of FCs as they synchronously engulf the germline cells [6,7] (Fig 1a). Investigation into the underlying genetic factors has identified important roles for effector caspase *Dcp-1* [8–10] and autophagy [11–14], indicating that the germline is eliminated by a combination of apoptosis and autophagic mechanisms. Further, FCs express the phagocytic receptor Draper (Drpr) on their cell surfaces which recognizes and internalizes the dying germline, leading to the subsequent activation of the JNK signaling pathway to promote resolution of clearance [6,7,15].

**Fig 1.**
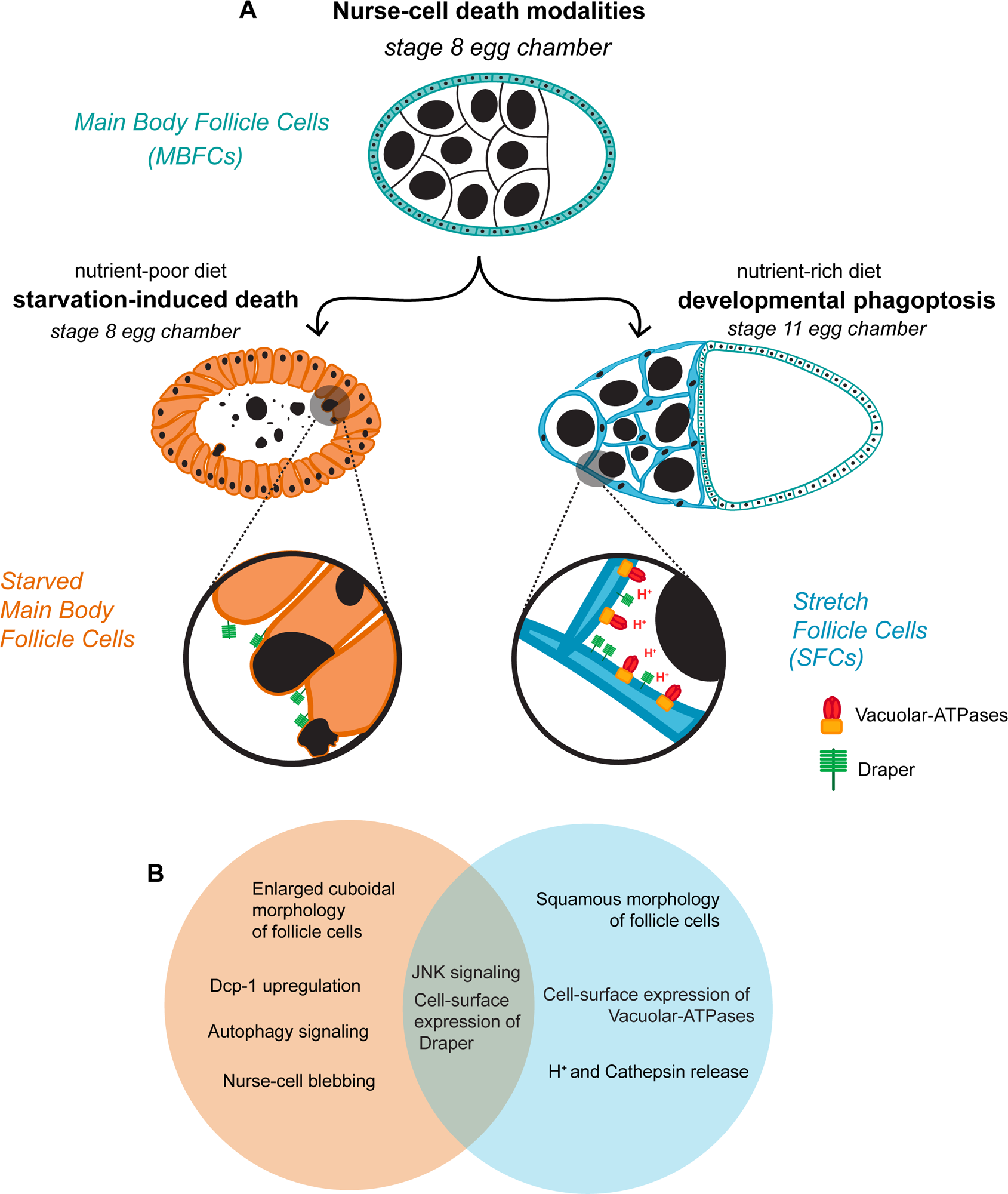
Nurse cell (NC) clearance is regulated apoptotically or non-apoptotically depending on environmental cues. (A) Schematic illustrating the different fates of egg chambers. A stage egg 8 chamber is completely surrounded by main body follicle cells (shown in teal). When the milieu is optimal and sufficient dietary protein is available, oogenesis proceeds through 14 stages. A subset of the main body follicle cells (MBFCs) differentiates into stretch follicle cells (SFCs, blue) and express the phagocytic receptor Draper and the proton pump Vacuolar-ATPase on their plasma membrane, ultimately acidifying the germline sister Nurse Cells (NCs, black nuclei) externally to clear them. In contrast, when the organism is deprived of nutrition, MBFCs enlarge in place (orange) and engulf the NCs, also by expressing Draper on their surfaces. In both death modalities, FCs clear NCs, albeit using different morphological configurations and biochemical pathways, providing a unique opportunity to study diverse cell death modalities in the same system (B) Venn diagram illustrating that developmental phagoptosis (blue) and starvation-induced death of NCs (orange) share key features such as the expression of Draper on the surface of the follicle cells and the activation of JNK signaling.

When the milieu is optimal and any physiological insults are overcome, oogenesis proceeds through 14 stages. A marked difference in egg chamber morphology is observed as it enters the vitellogenic phase in mid-oogenesis. A majority of the FCs migrate to encapsulate the oocyte, referred to as Main Body FCs (MBFCs), and a subset of them flatten and elongate to become Stretch Follicle Cells (SFCs) that migrate and extend into the spaces between the NCs [16] (Fig 1a). Stage 11 is defined by nurse cell “dumping”, in which maternal mRNAs, proteins and organelles from the NCs are deposited into the transcriptionally quiescent oocyte via ring canals, leaving behind the NC nuclei and a small amount of cytoplasm [3]. By stage 13, the NC nuclei begin to be eliminated non-autonomously and asynchronously by SFCs, in a process morphologically and physiologically distinct from starvation-induced NC clearance [17]. At the close of oogenesis at stage 14, all the NCs are cleared away, leaving only the fully developed oocyte. Genetic ablation of SFCs revealed that SFCs were required for NC dumping, death, and clearance [18], while disruption of caspases and autophagy-related genes failed to prevent NC death, indicating that developmental NC death occurs through non-apoptotic and non-autophagic mechanisms [18,19]. However, the usual suspects in engulfment - *drpr*, *Ced-12*, and the JNK signaling pathway, were found to be required for NC death and clearance, indicating that the phagocytosis machinery is deployed by SFCs [18]. Subsequently, the NCs are externally acidified by SFCs targeting their lysosomal machinery, specifically Vacuolar-ATPases (V-ATPases), to their plasma membranes [20]. This form of cell death, where the phagocytosis apparatus of one cell is harnessed to kill another cell, which would otherwise continue to live, is known as phagoptosis [21,22]. Recent findings on the molecular mechanisms underlying both phagoptosis and starvation-induced death modalities have been reviewed more comprehensively elsewhere [1,7].

In both of these forms of nurse cell death and clearance, epithelial follicle cells are recruited for NC elimination, acting as non-professional, tissue-resident phagocytes, albeit in distinct morphological configurations (Fig 1A). While there are differences in genes required in the NCs for their death, the two death regimes share some components, specifically in the engulfment machinery acting in FCs, indicating that clearance requires the preferential activation or repression of specific sub-routines of death-associated signaling cascades (Fig 1B). The follicle cells could also be primed differentially by distinct external cues, arising both from NCs, in the form of “find me” and “eat me” signals [23], as well as from the extracellular environ, shaped in part, by the nutrient sensing pathway. How follicle cells in mid-oogenesis are able to make context-driven decisions to steer the fates of nurse cells, and of themselves, remains to be determined.

To determine how follicle cells are deployed differently during the two types of NC death, we took a multimodal approach by capturing the “translatome” and “secretome” of follicle cells in these diverse contexts. We performed Translational Ribosome Affinity Purification with RNA-sequencing (TRAP-seq), in addition to proximity labeling (HRP-KDEL) followed by Liquid Chromatography tandem mass-spectrometry (LC-MS/MS) of MBFCs under fed and starved conditions, as well as after their transition to SFCs, to identify the pathways that differentially promote distinct nurse cell clearance routines. Further, we compared the translational profiles of fed MBFCs and fed SFCs to their transcriptional profiles obtained by integrating two previously published single-cell RNAseq atlases of the fly ovaries [24,25] to identify genes that are exclusively under translational regulation. Our analyses and validation identified several key genes and pathways that play vital roles during the making of a healthy egg, ranging from ensuring structural integrity and regulating metabolic pathways, to key signaling molecules. In summary, we have generated a multidimensional portrait of fly ovarian follicle cells with which we dissect and discern their complex, context-dependent behavior during NC death and clearance.

## Results

### Establishing the translatome, secretome, and transcriptome of MBFCs and SFCs

To capture the translatome or the set of all mRNAs undergoing translation in a cell, we performed TRAP-seq [26,27], by expressing GFP-tagged large ribosomal subunit 10a in follicle cells using the GAL4/UAS system [28,29]. We then performed immunoprecipitation to capture ribosome-bound mRNAs and subsequently sequenced and analyzed the transcripts. We used the follicle cell line GR1-GAL4 to express RpL10a-GFP in MBFCs (GR1>*RpL10a*-GFP) and PG150-GAL4 to express RpL10a-GFP in SFCs (PG150>*RpL10a*-GFP) (Fig 2A, S1 Fig). We generated biological triplicates corresponding to experimental groups of fed MBFCs, fed SFCs, and starved MBFCs, with each replicate averaging over 49 million reads (Fig 2B). Analysis with Salmon and DEseq2 identified 1133 differentially translated genes (FDR Adjusted p<0.05) associated with phagoptotic clearance of nurse cells by comparing fed SFC samples to the fed MBFC baseline. Similarly, comparison of starved MBFC samples to fed MBFCs identified 99 differentially translated genes (FDR Adjusted p<0.05). The functional roles played by these candidates is described in detail in subsequent sections.

**Fig 2.**
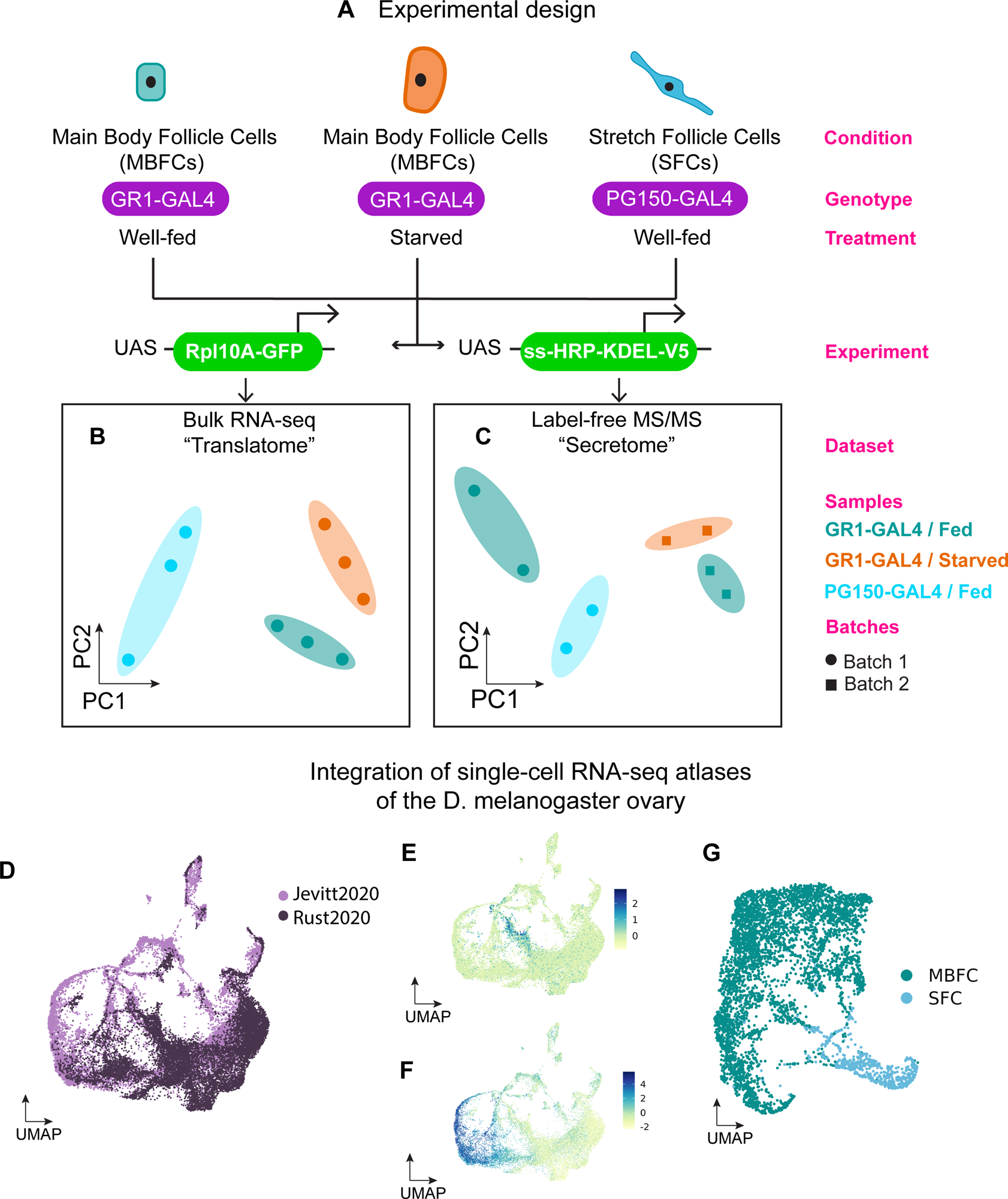
Establishing the translatome, secretome, and transcriptome of MBFCs and SFCs. (A) Schematic illustrating the experimental design for identification of the translatome and secretome from 3 conditions/cell types. *GAL4* drivers *GR1* and *PG150* were used to drive expression of *Rpl10a-GFP* in MBFCs (fed or protein-starved) and SFCs. The resulting GFP-conjugated ribosomes were immunoprecipitated and the mRNAs bound to the ribosomes were sequenced to produce the translatome. Similarly, to generate the secretome, *GR1* and *PG150* were used to drive the expression of *ss-HRP-KDEL-V5* in MBFCs and SFCs. HRP-KDEL expressed in cells localizes to the endoplasmic reticulum and biotinylated when incubated with biotin-phenol and H_2_O_2_. The tagged proteins were subsequently isolated using streptavidin beads and analyzed via LC-MS/MS to produce the secretome. (B) PCA Dimension reduction of aggregated gene-level transcript counts in translatome bioreplicates. (C) PCA of peptide intensity values of secretome bioreplicates. (D) UMAP embedding of Seurat RPCA integrated cells from publicly available single-cell RNA-seq atlases of the *Drosophila melanogaster* ovaries [24,25]. Cells are colored by the dataset of origin. (E) UMAP embedding of the integrated ovary datasets colored by SFC module score. To identify cells most likely to be stretch follicle cells in the integrated dataset, an aggregate score comprising the expression values of canonical SFC markers used in the published individual datasets were computed and projected on to the UMAP. (F) UMAP embedding of the integrated ovary datasets colored by mid/late MBFC module score. To identify cells most likely to be mid/late MBFCs in the integrated dataset, an aggregate score comprising the expression values of canonical mid/late MBFC markers used in the published individual datasets were computed and projected on to the UMAP. (G) Cells with a high SFC score (>1) and high mid/late MBFC score (>2x) were extracted from the integrated dataset and subject to reclustering. Updated PCA and UMAP embeddings were computed for the MFBC and SFC subset and clusters containing MBFCs and SFCs were identified after normalizing read counts in the new subset.

To establish the secretome or the set of all trans-membrane and secreted proteins of the follicle cells, we took a tissue and organelle-specific proteomics approach by combining the GAL4/UAS system with proximity-dependent protein labeling, followed by LC-MS/MS. Proteins fated to be secreted or trans-membrane are trafficked through the endoplasmic reticulum (ER) and Golgi network. To isolate and identify these proteins, we biochemically tagged and enriched proteins in the ER by expressing a genetically tagged fusion construct downstream of UAS, carrying horseradish peroxidase (HRP) fused to the ER-retention signal KDEL, along with secretion signal IgK leader sequence (ss) and the V5 epitope [30]. We expressed ss-HRP-KDEL-V5 in follicle cells, where it localizes to the ER. Like the translatome, to achieve MBFC-specific expression, we used GR1-Gal4 (GR1>ss-HRP-KDEL-V5) and to achieve SFC-specific expression, we used PG150-Gal4 (PG150>ss-HRP-KDEL-V5) (Fig 2A, Figure A,B in S2 Fig). Next, we exposed dissected ovaries to biotin-phenol substrate and a brief pulse of H_2_O_2,_ which catalyzes the biotinylation of the proteins being trafficked through the ER. We confirmed that the biotinylation is ER-specific with α-V5 and streptavidin-488 immunostaining and additionally verified that the biotinylation is contingent on the availability of the biotin-phenol substrate and H_2_O_2,_ (Figure C-H in S2 Fig). We then performed streptavidin enrichment followed by on-bead trypsin digestion and LC-MS/MS on the samples in two batches. Two replicates of fed MBFC samples were produced in each batch as a baseline to compare to the two replicates each of fed SFCs and starved MBFCs that were generated in separate batches (Fig 2C). Stringent database (MaxQuant) based spectral searching identified ∼2200 unique proteins in batch 1 samples, and ∼1800 in batch 2 samples. After removing decoys (reversed protein sequences used to gauge the false discovery rate) and potential contaminants (e.g. trypsin), we obtained a high confidence set of 40 differentially abundant proteins in the phagoptosis comparison and 14 proteins in the starvation-induced clearance comparison (FDR Adjusted p<0.05) (Figure I-J in S2 Fig).

Next, we leveraged two previously published single-cell RNAseq atlases of the fly ovary [24,25] to obtain complete transcriptomic profiles of MBFCs and SFCs, against which we compared our findings from the translatome and the secretome, as well identified gene regulatory programs with finer resolution. The Jevitt et al. atlas [25] comprises cells from egg chambers across all developmental time points. However, the Rust et al. atlas [24] does not include vitellogenic egg chambers but delineates precursors to SFCs. Therefore, we integrated the two datasets using Seurat Reciprocal PCA and obtained a reclustered subset of cells that include putative MBFCs and SFCs across the developmental continuum (Fig 2D).

To identify MBFC and SFC populations comparable to the translatome and secretome, we obtained a cell identity module score using Seurat *AddModuleScore* based on the expression of canonical MBFC and SFC markers, such as *Yp1* and *cv-2* respectively (Figure A-E in S3 Fig) [31]. For the scope of this study, we bisected the reclustered subset broadly into MBFCs and SFCs and identified differentially expressed genes between the two groups and confirmed their concordance with findings from the original publications (Fig 2E-2G). We used the direction and magnitude of Log_2_ fold change (LFC) values from this comparison (SFCs / MBFCs) to contrast findings from the translatome and the secretome.

### Distinct genes contribute to overlaps in gene ontology (GO) terms enriched in phagoptosis and starvation-death of NCs

To identify shared and disjointed pathways between phagoptosis (fed SFCs vs. fed MBFCs) and starvation-death of NCs (starved MBFCs vs. fed MBFCs), we compared the LFC estimates from both the translatome and the secretome (Fig 3A-3B). For the sets of candidates identified in each dataset, we performed GO-term and KEGG pathway enrichment analysis [32]. We identified 5 congruently upregulated genes in the translatomes of SFCs and starved MBFCs (Fig 3C) involved in cytoskeletal remodeling (GO:0030239, GO:0061640, GO:0031032) and 49 congruently downregulated genes, a majority of which are associated with vitelline membrane and chorion deposition (GO:0007304, GO:0030703) (Fig 3D). We also identified other genes associated with the same GO-terms but individual genes were unique to either SFCs or starved MBFCs, highlighting the molecular differences underlying the similar processes that give rise to diverse phenotypic outcomes. The 415 genes uniquely upregulated in the SFC translatome were enriched for inositol and tyrosine metabolism (GO:0006020, GO:0006570), defense response (GO:0006952), and organic acid catabolism (GO:0016054). Accordant with our expectations, we found glucose homeostasis (GO:0042593), response to starvation (GO:0042594), and lipid metabolism (GO:0008610) over-represented exclusively in the starved MBFC translatome. Notably, response to wounding (GO:0009611) was also over-represented in this group (S1 Table).

**Fig 3.**
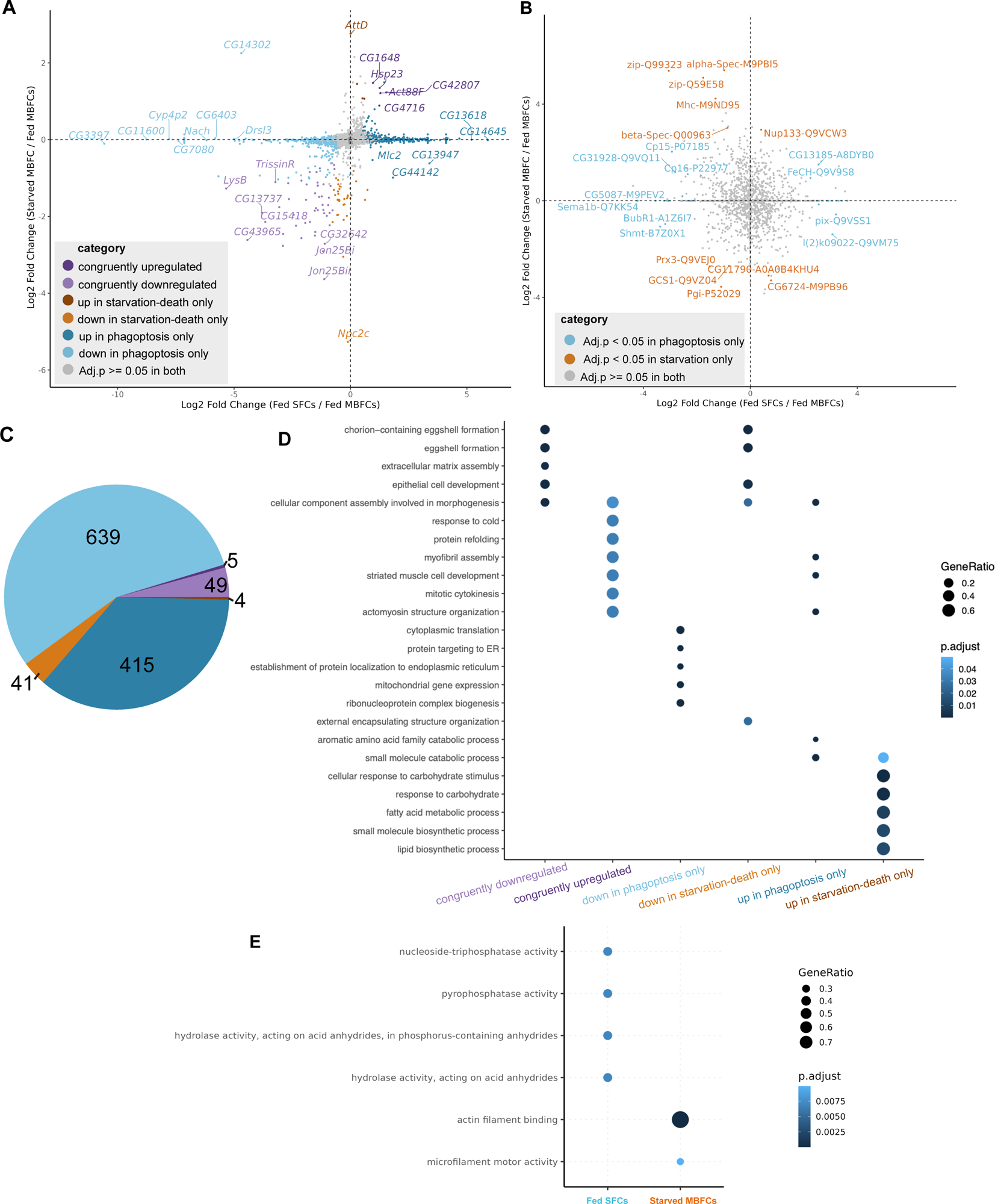
Functional enrichment of candidates identified in the translatome and secretome reveals pathways common and unique to phagoptosis and starvation-death of NCs. (A) Scatterplot of differentially translated gene candidates in phagoptosis (Fed SFCs / Fed MBFCs – x-axis) and starvation-death (Starved MBFCs / Fed MBFCs – y-axis) obtained using DESeq2. LFC differentials and FDR adjusted p-values from both analyses were compared to find genes congruently or uniquely regulated in both death modalities. Quadrants 1 and 3 indicate genes congruently up and down-regulated in phagoptosis and starvation-death respectively. Congruency was determined if the direction of LFC was the same and if the adjusted p-value was less than 0.05 in both. (B) Scatterplot of differentially abundant candidates in phagoptosis and starvation-death in the secretome. Proteins common to both death modalities were not found. UniProt identifiers along with gene symbols of detected proteins are specified. (C) Counts of genes regulated congruently and uniquely in phagoptosis and starvation-death of NCs in the translatome. Figure color legend same as in (A). (D) Dot plot of GO-terms enriched in the categories of differentially translated genes identified in (A and C). (E) Dot plot of GO-terms enriched in the secretome of fed SFCs and starved MBFCs.

While the candidates identified in the secretome were different from those identified in the translatome in both phagoptosis and starvation, we observed that the GO-terms over-represented in the corresponding comparisons were equivalent. For example, we found actin and microtubule associated proteins over-represented in the starved MBFC secretome (GO:0003779, GO:0008017, GO:0005200). In the SFC secretome, we found enrichment of proteins involved in the hydrolysis of acid anhydrides (GO:0016817) (Fig 3E). We did not find any congruity in the differentially abundant proteins identified in phagoptosis and starvation-death in the secretome.

We also compared the LFC values of differentially expressed genes between SFCs and MBFCs identified from the translatome and the integrated scRNAseq transcriptome. After filtering out genes with adjusted p value >= 0.05 in both datasets, we found 18 genes concordantly upregulated and 45 genes concordantly downregulated in SFCs (Figure F-H in S3 Fig). A majority of these genes were involved in cytoskeletal remodeling and cell morphogenesis or in eggshell deposition. Overall, we found only *Vm32E* and chorion associated gene *CG12398* downregulated in SFCs across all 3 datasets (S2 Table).

### SFCs and starved MBFCs congruently downregulate eggshell deposition

The *Drosophila* eggshell is a multi-layered extra-cellular matrix structure proximal to the oocyte, helping maintain egg chamber structural integrity. In mid and late oogenesis, MBFCs and oocytes secrete sequentially, several proteins that comprise the vitelline membrane and the several layers that comprise the chorion [33,34]. Genes required for optimal eggshell assembly such as *dec-1*, *Cp36*, and *Vm26Ab* were identified through screening of female sterile (fs) mutants [35–37]. In agreement with these findings, GO-term enrichment of our multi-modal dataset revealed that several genes involved in various eggshell deposition processes were downregulated in SFCs at the transcriptome, translatome, and at the secretome level, confirming that the datasets are capturing the appropriate tissues, as well as confirming that eggshell deposition is a MBFC-specific trait. Further, we used the GLAD matrisome gene list [38] to annotate our multi-modal dataset and comprehensively identify patterns of regulation in specific structural and functional sub-classes of the matrisome. We found that vitelline membrane-associated genes *Vm26Aa*, *Vm26Ab*, *Vm26Ac*, *Vm32E*, *Vm34Ca*, and *Vml* were all congruently downregulated in fed SFCs compared to the fed MBFC baseline in both the translatome and the integrated scRNAseq transcriptomes. However, only *Vm32E* was significantly downregulated in SFCs in the secretome, while the directionality of fold change of other Vm genes in the secretome still remained concordant with the other datasets. Similarly, chorion-related genes *Cp15*, *Cp16*, *Cp19*, *Cp36*, *Cp18*, and *Cp38* were significantly downregulated in SFCs in the translatome, while only *Cp15* and *Cp16* were significantly downregulated in SFCs at the secretome level. Interestingly, we observed Semaphorin 1b (*Sema1b*) and BMP ligand *gbb* were also downregulated by SFCs in the secretome, however this effect was not observed in the other datasets (S2 Table).

Previous studies have reported that starvation stress deters egg chambers from entering vitellogenesis [39]. In concordance, our analyses show that protein-deprived MBFCs significantly downregulate the translation of *Vm32E*, *Vml*, and all chorion-related genes. In addition to these apical-matrix associated genes, we found other extracellular matrix (ECM)-affiliated genes such as ECM-regulator *PH4alphaPV* and immune-regulated glycoprotein *I(2)34Fc* to be downregulated in the translatome of starved MBFCs. However, we did not observe a similar trend in their corresponding protein products in the secretome.

### Cytoskeletal dynamics in SFCs and starved MBFCs

During stages 9 and 10, the egg chamber further elongates and maintains an anterior-posterior axis while the oocyte grows tremendously to occupy over half the volume of the egg chamber. Keeping up with the increasing surface area of the egg chamber, anterior follicle cells elongate and flatten, acquiring the characteristic squamous morphology, cellular patterning, and identity of SFCs. It was previously [40] reported that downregulation of *Br* expression mediated by ecdysone and JAK/STAT pathways is essential for SFC flattening [40]. Several additional genes involved in the cuboidal-to-squamous transition of follicle cells, such as *Tm1*, *Rab11*, and *Zip* were also reported [40]. In contrast, during starvation-induced clearance, MBFCs undergo cytoskeletal rearrangements to enlarge in place, and the GTPase Rac1 has been previously reported to be required for the cytoskeletal rearrangements during phagocytosis. We previously reported that *Rac1* is required in starved MBFCs to promote the enlargement of FCs and engulfment of apoptotic nurse cells [6]. Rac1 is also required in SFCs even though acidification, but not engulfment of nurse cells has been observed in phagoptosis [18]. Several key questions related to cytoskeletal dynamics remain unanswered. Namely, how are follicle cells able to assume two different morphological configurations in phagoptosis and starvation-induced clearance of nurse cells? Are the same morphogenetic events required to surround and internalize nurse cell targets in both death paradigms? What controls SFC density and patterning along the anterior-posterior axis?

Towards this goal, we identified 26 cytoskeleton-associated genes upregulated in the translatomes of SFCs. In concordance with the SFC transcriptome, we found *Tm1* upregulated in the SFC translatome, along with several other calcium-dependent regulators of the cytoskeleton, such as *Tm2*, *up*, *Actn*, *Mlc1*, *Mlc2*, *bt*, *TpnC41C*, *Scp1*, *cue*, and *didum* (S2 Table). From the SFC secretome, we found a potential role for *armi* in late oogenesis, which closely associates with the microtubule cytoskeleton in egg chambers during early and mid-oogenesis, influencing egg chamber axial polarity during these stages [41]. Of note, we found actin encoding *Act88F* to be upregulated in both SFC and starved MBFC translatomes. In addition, *Act79B* upregulation was exclusive to the SFC translatome. Because Ca^2+^ signaling plays a vital role in actin polymerization and phagocytic cup formation in other systems [42], we characterized the consequences of disrupting *TpnC41C*, *Scp1*, and *cue* by RNAi, using the GR1 driver. All 3 candidates demonstrated a persisting NC nuclei (PNCN) phenotype indicating their requirement in phagoptotic NC clearance. In addition, we performed RNAi of other cytoskeleton regulators and observed a weak PNCN phenotype (Fig 4A-4C).

**Fig 4.**
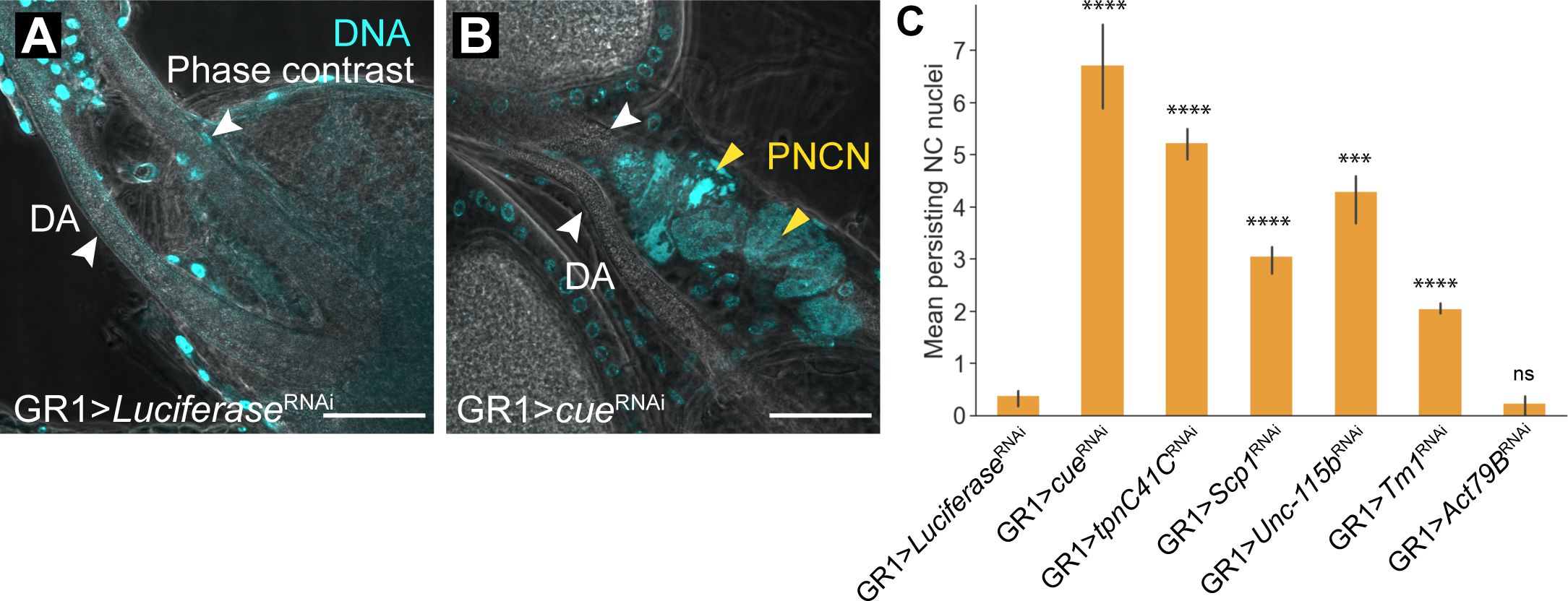
RNAi knockdown of Ca2+ associated cytoskeleton regulators leads to developmental NC clearance defects. (A-B) Representative images of anterior regions of stage 14 egg chambers from control (GR1 > *Luciferase* RNAi) and mutant (GR1 > *cue* RNAi) flies. DNA is labelled with DAPI (cyan). Stage 14 egg chambers are distinguished by two fully developed dorsal appendages (DA), indicated by white arrows. Persisting NC nuclei (PNCN) indicating a disruption to phagoptosis resulting from RNAi are marked by yellow arrows. Scale bars = 50µ. (C) Quantification of PNCN in stage 14 egg chambers. GR1 > *cue* RNAi demonstrates a strong PNCN phenotype, while the other mutants demonstrate a moderate to weak PNCN phenotype. At least 10 flies (20 ovaries) were used for each genotype. (*** p <= 5e-04, **** p <= 5e-05, ns p>0.05, one-sided independent t-test)

### Regulation of lysosomal machinery during phagoptosis of nurse cells

We previously reported that the lysosomal machinery, in particular V-ATPases, was required in SFCs for the acidification of nurse cells [20]. Our analysis of the integrated scRNAseq dataset identified 15 genes associated with V-ATPases that were upregulated by SFCs at less than 1.5x LFC differential (S2 Table). By contrast, we found only *Vha44* to be upregulated by SFCs in the translatome in concordance with the transcriptome. Paradoxically, we found *Vha68-1* to be upregulated by SFCs in the translatome, while *Vha68-2*, another gene isoform that codes for the same V-ATPase subunit (V1 catalytic domain, subunit A) to be upregulated in SFCs in the transcriptome. At the secretome level, none of the V-ATPase protein products showed a significant differential even though peptides of various subunits were identified. In addition to V-ATPases, we identified 18 lysosome-associated genes enriched in the SFC translatome. These include a transmembrane lysophospholipase *sws*, *g* (*garnet*) which encodes a subunit of lysosomal adapter protein complex AP-3, and *psidin*, which also affects lamellipodial dynamics in border cells, as well as initiates humoral immune response [43,44]. Likewise, the SFC secretome included 3 lysosomal proteins Plekhm1, FeCH, and CG43693. Notably, mutations in the human ortholog of Plekhm1 have been implicated in autosomal recessive osteopetrosis [45], which results from improper lysosomal acidification and resorption of the extracellular bone matrix [46], a mechanism reminiscent of SFC acidification of nurse cells. Interestingly, the SFC secretome was also enriched for four ATPases Ns1 (Q8MT06), Ns2 (Q7JXU4), pix (Q9VSS1), and CG13185 (A8DYB0), which are associated with hydrolysis of acid anhydrides, a reaction which results in the formation of carboxylic acids. Intrigued by this finding, we examined the SFC translatome further, finding 11 upregulated genes that are involved in carboxylic acid metabolism (GO:0016054, GO: 0019752). These included *hgo, Faa, Gabat, CG1440, CG3699, CG1461, Hn, Spat, Hpd, SERCA,* and *GstZ2*.

### The role of innate immune signaling in NC death and clearance

The innate immune response in *Drosophila* relies on two highly conserved NF-κB pathways - Toll and Immune Deficiency (Imd). When the organism is immunologically challenged, a class of circulating secreted molecules called peptidoglycan recognition proteins (PGRPs) recognize and bind pathogen associated molecular patterns (PAMPs), such as Lys-type or DAG-type peptidoglycans or β-glucan which are expressed on the pathogen [47,48]. Depending on the infecting pathogen, Toll or Imd signaling is triggered preferentially.

Toll is typically activated upon infection by gram-positive bacteria or fungi, which initiates a serine protease cascade in host cells, whereby the zymogen Spatzle (Spz) is activated by proteolysis, after which it binds to the membrane-bound Toll receptor [49]. Spatzle Processing Enzyme (SPE), a serine protease that cleaves Spz to activate Toll, was later identified and characterized [50] The intracellular Toll/Interleukin-1 receptor domain of the Toll receptor then binds to the adapter protein MyD88, which subsequently forms a complex with kinase Pelle and adapter protein Tube, leading to the phosphorylation and degradation of the *Drosophila* IκB factor Cactus [51,52]. This allows the translocation of the NF-κB family transcription factors Dif and Dorsal, ultimately activating the expression of genes encoding several antimicrobial peptides (AMPs) such as *Drosomycin* (*Drs*) and *AttacinA* (*AttA*) to combat infection [53]. When the cascade is triggered, the AMPs produced are embedded in the cell envelopes of pathogens, subsequently destabilizing and killing the pathogen [53]. Loss of *SPE* has been shown to impair *Drs* induction in response to microbial infection [50]. The Spz-Toll pathway also plays a role in development, and these pathways have also been shown to promote neurodegeneration, so their roles extend beyond pathogen response.

Our analysis of the translatome revealed that SFCs upregulated not only the upstream peptidoglycan recognition molecules *PGRP-LC* (1x LFC) and *PGRP-SD* (1.5x LFC), but also the protease *SPE* (1x LFC) and AMP *Drs* (2.8x LFC). Furthermore, we observed several other known and predicted NF-κB/Toll regulators and AMP genes upregulated in the SFC translatome (Fig 5A). To investigate the role of NF-κB/Toll signaling components in phagoptosis, we performed RNAi against *PGRP-SD, PGRP-LC*, *SPE*, *Drs*, and *Drsl4* and overexpressed *Listericin* using the GR1 driver. All of the transgenic lines, except *SPE* demonstrated a moderate or weak PNCN phenotype (Fig 5B). Surprisingly, we found that loss of *SPE* caused significant, widespread egg chamber degeneration in mid-oogenesis, resulting in near-complete loss of late-stage vitellogenic egg chambers. Almost all ovarioles in GR1 > *SPE* RNAi mutants demonstrated an intact germarium and healthy early-stage egg chambers which began degenerating around stage 6 (Fig 5C-5D). Antibody staining against Discs large (Dlg), a scaffolding protein that stains plasma membranes, showed that unlike starvation-induced degeneration of wildtype egg chambers in mid-oogenesis, GR1 > *SPE* RNAi egg chambers neither demonstrated enlargement of MBFCs nor engulfment and clearance of NCs. MBFCs in GR1 > *SPE* RNAi egg chambers instead were partially or completely missing in younger egg chambers, indicating that MBFCs are affected prior to NC degradation due to loss of *SPE*. Further, NC chromatin in GR1 > *SPE* RNAi mutants appeared highly condensed, with some NC nuclei also undergoing fragmentation. Of note, the anterior-posterior polarity of egg chambers was also affected, with some dying NCs encroaching into the posterior end, displacing the oocyte. Several degenerating egg chambers did not show any detectable oocyte (Fig 5E-5G). While our experiments show that *SPE* is required for proper oogenesis, it is unknown whether the canonical Toll pathway is activated in follicle cells. While our bioinformatics analysis identified multiple components of the Toll pathway, including AMPs upregulated in the SFC translatome it remains to be determined if *SPE* acts in the canonical pathway to cleave *Spz* and if the AMPs upregulated have an immunogenic role in phagoptosis of NCs.

**Fig 5.**
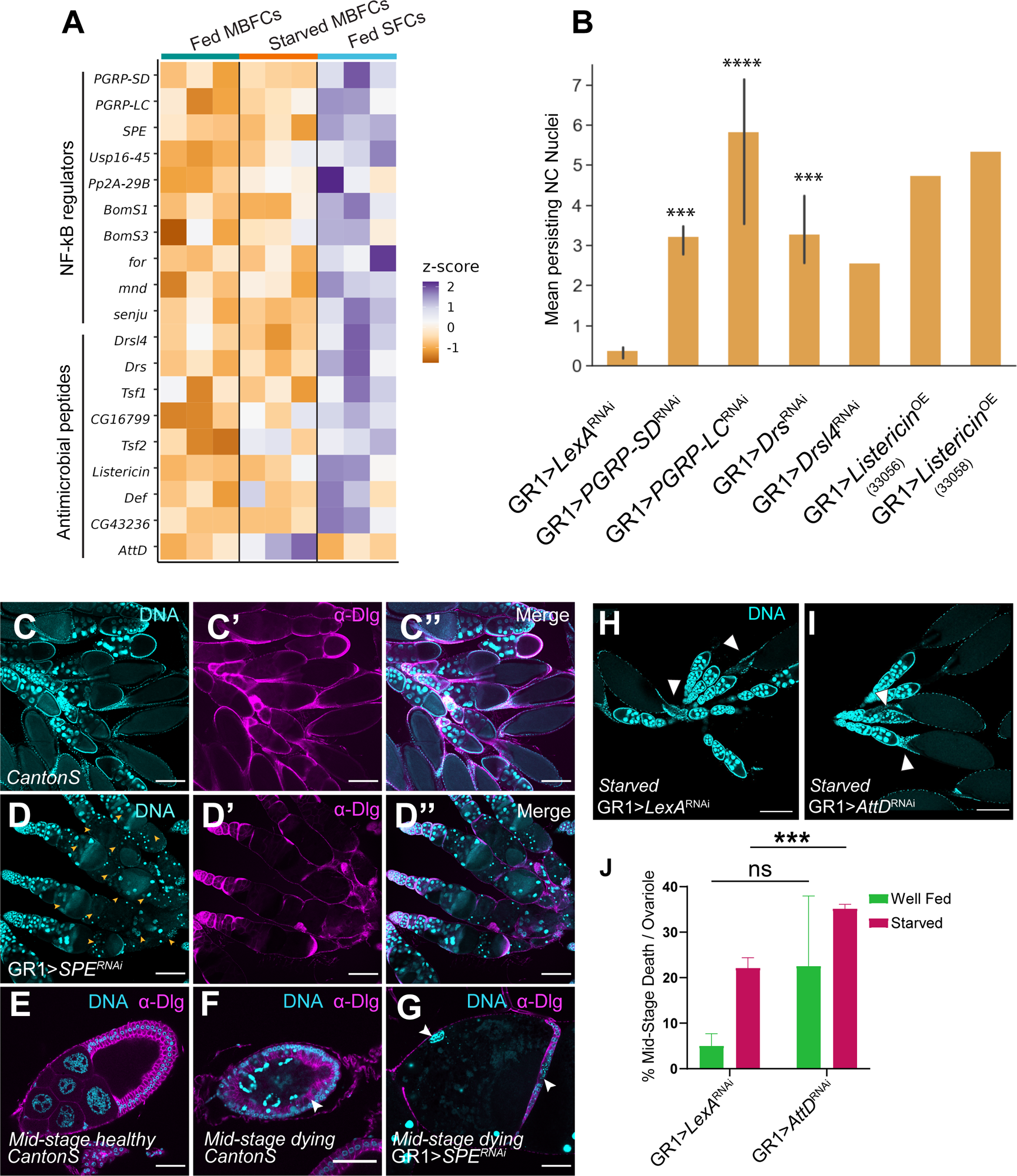
The role of innate immune signaling in NC clearance. (A) Heatmap of scaled read counts of NF-κB regulators and antimicrobial peptide genes upregulated by fed SFCs or starved MBFCs in the translatome. (B) Quantification of PNCN in stage 14 egg chambers from NF-κB or AMP knockdowns. (*** p <= 5e-04, **** p <= 5e-05, ns p>0.05, one-sided independent t-test). Significance levels are indicated for samples that have at least 3 replicates. (C) Egg chambers from wild-type fly ovaries stained with DAPI (cyan) and anti-Dlg (magenta) to label FC membranes. Scale bars= 200µ. (D) Egg chambers from *GR1 > SPE RNAi* fly ovaries stained with DAPI and anti-Dlg. *GR1 > SPE RNAi* ovarioles contain intact germaria and healthy early egg chambers but show wide-spread degeneration beginning around stage 6. Scale bars= 200µ. (E-G) Images of individual healthy and mid-stage dying egg chambers from wild-type ovaries and a mid-stage dying egg chamber from *GR1 > SPE RNAi* stained with DAPI and anti-Dlg. The *GR1 > SPE RNAi* degenerating egg chamber is lacking most of the MBFC layer, indicated by white arrows. NC DNA in *GR1 > SPE RNAi* is highly condensed and few are fragmented. Anterior-posterior polarity of the egg chamber is lost, with NCs extending into the posterior end, displacing the oocyte. Scale bars= 50µ. (H) *GR1 >LexA RNAi* starved control shows sporadically degenerating egg chambers, Scale bars= 200µ. (I) *GR1 >AttD RNAi* starved shows increased degeneration of egg chambers Scale bars= 200µ. (J) Quantitative analysis of midstage degenerating egg chambers with one-way ANOVA.

In contrast to *Drs* upregulation in phagoptosis, we found AMP gene *Attacin D (AttD)* to be upregulated in the starved MBFC translatome. This was particularly intriguing as some Attacin genes encode glycine-rich AMPs, typically expressed in response to Gram-negative bacterial liposaccharides, regulated by the Imd pathway. Unlike the Toll pathway, which leads to the nuclear localization of Dif and Dorsal, the Imd pathway leads to the nuclear translocation of NF-κB transcription factor *Relish (Rel),* which results from the phosphorylation of the N-terminus, as well as cleavage of C-terminus of Rel [48,54,55]. We wondered whether *AttD* functions as an AMP in starved MBFCs to destabilize and promote apoptosis of NCs and if loss of *AttD* would attenuate egg chamber degeneration in response to starvation. Surprisingly, we found that RNAi knockdown of *AttD* in starved MBFCs resulted in an increase in the number of mid-oogenesis deaths per ovariole. Additionally, our analyses showed that *AttD* RNAi in follicle cells did not cause increased mid-stage death in well-fed flies, indicating that the transgenic construct is not sufficiently lethal to the egg chamber by itself (Figure 5H-5J). This indicates an alternative role for *AttD* during starvation stress. One possibility is that *AttD* from MBFCs of degenerating egg chambers functions as negative feedback, preventing or limiting apoptosis to preserve younger egg chambers. It was previously reported that overexpressing *imd* did not activate pro-apoptotic genes *hid* and *reaper* [56]. Further, activating the Imd pathway upregulated the inhibitor of apoptosis *Diap1* in the larval epidermis and fat body, indicating that Imd activation could serve multifarious functions. While *AttD* showed 2.8x LFC upregulation in the starved MBFC translatome, we did not find upstream regulators of Imd or other Imd-associated AMPs to indicate the activation of the canonical Imd pathway (S3 Table). Interestingly *PGRP-LC*, which is typically associated with Imd activation was instead found upregulated in fed SFCs along with Toll regulators.

### *in vivo* RNAi screening of candidates

In addition to previously described gene families, we performed *in vivo* validation for a number of candidates identified across the 3 datasets presented herein (Table 1). A majority of the candidates we elected to follow up with were known secreted or transmembrane proteins, in order to identify any key components that may be involved in cell-cell communication in both death modalities. We also identified other candidates involved in major *Drosophila* signaling pathways to be differentially regulated exclusively in phagoptosis (Fig 6A). Specifically, we found *Dad* (*Daughters against dpp*) a component of the TGF-β/BMP signaling pathway to be enriched in SFCs in both the integrated scRNAseq data and the translatome. Other genes in this pathway such as *cv-2* and *dpp* have been previously shown to be expressed in SFCs [16]. RNAi against *Dad* in follicle cells resulted in a weak PNCN phenotype, instead exhibiting a strong “dumpless” phenotype, in which NCs fail to dump their contents into the oocyte while the dorsal appendage continues to form (Fig 6B). We also observed several morphological defects, ranging from fused egg chambers and distended SFC membranes, to stunted dorsal appendages, indicating that *Dad* and the TGF-β/BMP pathway might also play a role in SFC differentiation and patterning in late-oogenesis. RNAi against several other candidates such as *Tm1*, *LBR* (Fig 6C), and *magu* also presented a dumpless phenotype (Table 1). Of all the candidates we tested, GR1 > *prosα3 (proteasome α3 subunit)* RNAi stage 14 egg chambers had the strongest persisting NC nuclei phenotype (Fig 6E, 6H), with some egg chambers retaining all 15 NC nuclei. In addition, this genotype also had egg chambers with several morphological defects, including stunted dorsal appendages, and increased mid-stage death even when fed a protein-rich diet. When starved, this genotype presented the same morphological defects and strong PNCN phenotype, in addition to increased midstage death. One of the most interesting mutant phenotypes we observed was in GR1 > *wat* RNAi, which presented abnormal NC nuclei, starting around stage 8. NC nuclei in GR1 > *wat* RNAi egg chambers had large areas devoid of DAPI staining, when compared to CantonS controls (Fig 6F, 6G). However, they did not appear to be pyknotic nor strongly persist in stage 14 egg chambers (Fig 6H). For all of the candidates tested, we qualitatively and quantitatively summarized mutant phenotypes, broadly categorized into 4 classes – NC clearance defects, NC dumping defects, egg chamber morphological and patterning defects, and excessive degeneration of egg chambers in mid-oogenesis (Fig 6H, Table 1).

**Fig 6.**
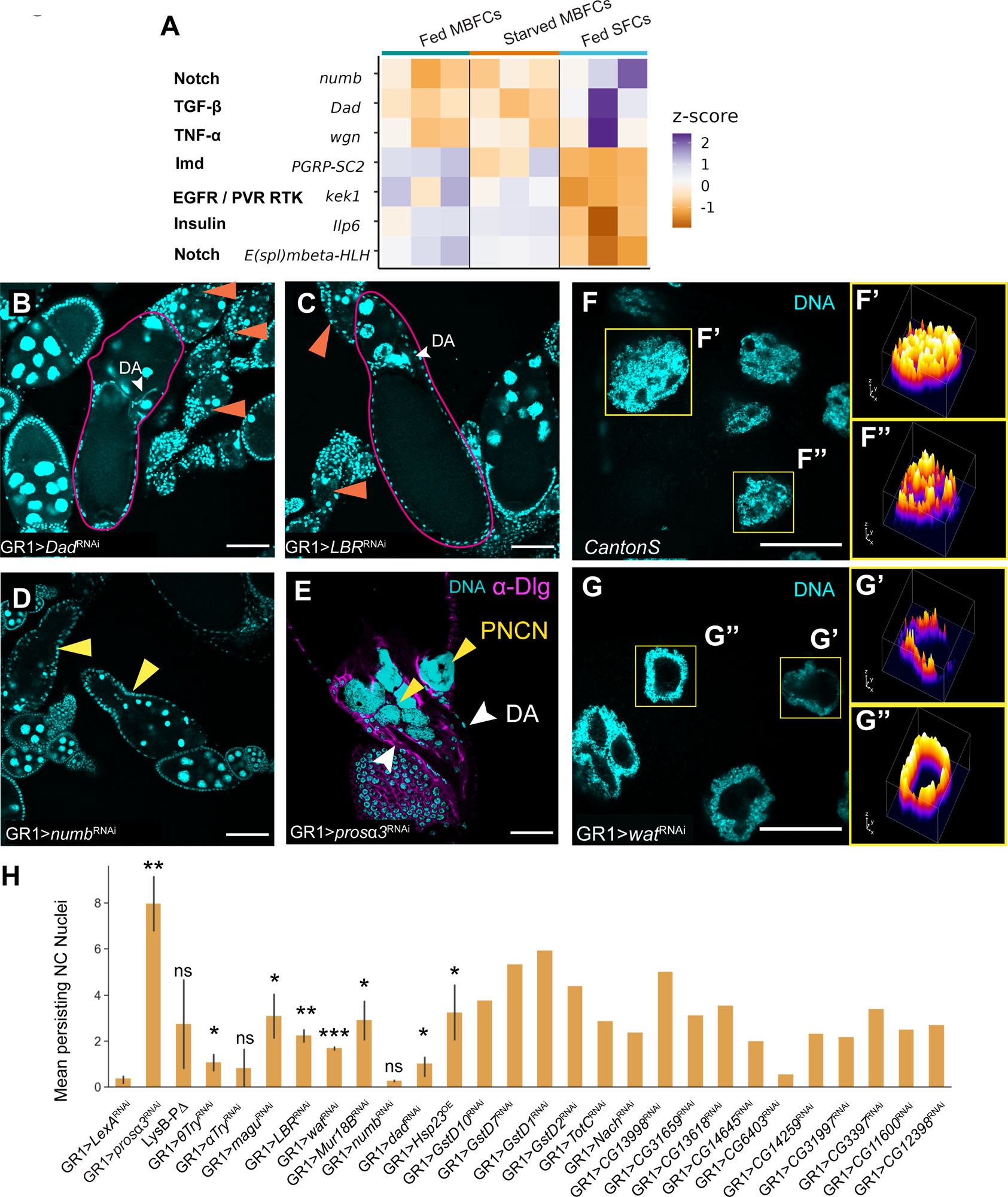
*in vivo* RNAi screening under fed condition. (A) Heatmap of scaled read counts of major signaling pathway genes that are significantly differentially regulated in SFCs. (B,C) *GR1 > Dad RNAi* and *GR1 > LBR RNAi* result in a “dumpless” phenotype (outlined in magenta), in addition to increased mid-stage death (orange arrows). Scale bar=100µ. (D) *GR1 > numb RNAi* results in mid-stage death accompanied by abnormal egg chamber morphology (yellow arrows). Scale bar=100µ. (E) *GR1 > prosα3 RNAi* ovaries have a strong PNCN phenotype. Scale bar=50µ. (F-G) GR1 > *wat* RNAi results in extensive NC nuclei vacuolization. (F) NC nuclei in a stage 9 egg chamber from CantonS ovary. Scale bar = 50µ. (F’, F’’) 3D surface plot of individual nuclei in stage 9 egg chamber from CantonS ovary after z-projection (max-intensity across stacks). Colors of peaks represent pixel intensity across z-stacks, with darker colors representing lower pixel intensity in the area (low DAPI staining) and lighter colors representing higher pixel intensity (robust DAPI staining). (G) NC nuclei in a stage 9 egg chamber from *GR1 > wat RNAi* ovary. Scale bar = 50µ. (G’, G’’) 3D surface plot of individual nuclei in stage 9 egg chamber from *GR1 > wat RNAi* ovary after z-projection (max-intensity across stacks) reveals increased vacuolization. (H) Quantification of PNCN in stage 14 egg chambers of candidates from Table 1. (* p < 5e-02, ** p <= 5e-03, *** p <= 5e-04, **** p <= 5e-05, ns p>0.05, one-sided independent t-test). Significance levels are indicated for samples that have at least 3 replicates.

**Table 1.** Summary of results from the *in vivo* validation of candidates identified from the translatome, secretome, and the integrated scRNA-seq datasets. For each candidate, the rationale for additional follow-up, RNAi or overexpression stocks used, summary of phenotypes observed in well-fed and starved experiments are listed. Stocks are RNAi lines unless otherwise indicated (*=OE, #=CRISPR/Cas9 deletion)

| Gene Symbol | Enriched in | Support | Gene-Ontology Annotation | Stock | Phenotype summary |
| --- | --- | --- | --- | --- | --- |
| **Nach** | fed MBFCs | translatome | Secreted/Transmem | 27262 | Weak PNCN |
| **to** | fed SFCs | translatome | Secreted/Transmem | 27646* | Moderate PNCN |
| **CG33470 & IMPPP** | fed SFCs | translatome | Secreted/Transmem | 28540 | Weak PNCN |
| **Listericin** | fed SFCs | translatome | Secreted/Transmem | 33058* | Moderate PNCN |
| **Listericin** | fed SFCs | translatome | Secreted/Transmem | 33056* | Moderate PNCN |
| **CG13998** | fed MBFCs | scRNAseq, translatome | Secreted/Transmem | 34599 | Moderate PNCN |
| **Drsl4** | fed SFCs | translatome | Secreted/Transmem | 57186 | Dumpleless, Weak PNCN |
| **CG31659** | fed MBFCs | translatome | Secreted/Transmem | 57203 | None |
| **CG13618** | fed SFCs | translatome | Secreted/Transmem | 57749 | Dumpleless, Weak PNCN |
| **CG14645** | fed SFCs | translatome | Secreted/Transmem | 60068 | Weak PNCN |
| **CG31999** | fed SFCs | translatome | Secreted/Transmem | 31587 | Weak PNCN, Dumpleless |
| **LysB-Pdelta** | fed MBFCs | translatome | Secreted/Transmem | [57]# | several Dumpleless |
| **TotC** | fed MBFCs | translatome | Secreted/Transmem | 62159 | Weak PNCN |
| **magu** | fed SFCs | scRNAseq, translatome | Secreted/Transmem | 51169 | Dumpleless |
| **magu** | fed SFCs | scRNAseq, translatome | Secreted/Transmem | 41641 | Dumpleless |
| **alphaTry** | fed MBFCs | translatome | Secreted/Transmem | 51824 | Dumpleless |
| **thetaTry** | fed MBFCs | translatome | Secreted/Transmem | 53362 | None |
| **Mur18B** | fed SFCs | translatome | Secreted/Transmem | 56957 | Dumpleless, constricted egg chambers |
| **CG14259** | fed SFCs | translatome | Secreted/Transmem | 57196 | Moderate PNCN, Dumpleless |
| **CG31997** | fed SFCs | translatome | Secreted/Transmem | 57517 | Moderate PNCN, Dumpleless, Shriveled unhealthy stage 14 egg chambers |
| **wat** | fed SFCs | translatome | Secreted/Transmem | 67801 | Dumpleless, abnormal nuclear morphology |
| **LBR** | fed SFCs | translatome, secretome - NS | Secreted/ Transmem | 53269 | Weak PNCN, severe Dumpless, midstage death |
| **Prip** | fed SFCs | translatome, secretome - NS | Secreted/ Transmem | 50695 | Moderate mid-stage death, Dumpless |
| **AttD** | starved MBFCs | translatome | Secreted/ Transmem | 55979 | None in fed, Higher mid-stage death in starved |
| **CG10163** | starved MBFCs | translatome - NS | Secreted/ Transmem | 32959 | Higher mid-stage death in fed |
| **CG31926** | starved MBFCs | translatome | Secreted/ Transmem | 64664 | None in fed, none in starved |
| **Lsp1beta** | starved MBFCs | translatome | Secreted/ Transmem | 27042 | None in fed, none in starved |
| **Drs** | fed SFCs | translatome | Secreted/ Transmem | 55391 | Weak PNCN, Dumpless |
| **CG17192** | starved MBFCs | translatome | Secreted/ Transmem | 56042 | None in fed, none in starved |
| **CG11600** | fed MBFCs | translatome | Metabolic | 66927 | None |
| **CG3397** | fed MBFCs | translatome | Metabolic | 44115 | Weak PNCN, Dumpless |
| **Cyp6a17** | fed SFCs | translatome | Metabolic | 33887 | None |
| **GstD10** | fed MBFCs | translatome | Metabolic | 64678 | Weak PNCN |
| **GstD7** | fed MBFCs | translatome | Metabolic | 65961 | Weak PNCN |
| **GstD1** | fed MBFCs | translatome | Metabolic | 77338 | Moderate PNCN, Dumpless |
| **GstD2** | fed MBFCs | translatome | Metabolic | 77361 | Weak PNCN, Dumpless |
| **CG12398** | fed MBFCs | scRNAseq, translatome, and secretome | Metabolic | 62837 | Weak PNCN |
| **Prosalpha3** | fed SFCs | scRNA & translatome, but enriched in Fed MBFCs in secretome | Metabolic | 77145 | Strong PNCN, stunted dorsal appendages, weak to moderate midstage death, severe arrested development, displaced and pyknotic NC nuclei, NCs in corpus luteum in fed. Severe midstage death and other previously listed phenotypes in starved |
| **CG6277** | starved MBFCs | translatome | Metabolic | 33958 | None in fed, none in starved |
| **CG6277** | starved MBFCs | translatome | Metabolic | 51030 | None in fed, none in starved |
| **CG17633** | starved MBFCs | translatome | Metabolic | 60043 | None in fed, none in starved |
| **CG6508** | starved MBFCs | translatome | Metabolic | 64953 | None in fed, none in starved |
| **Jon25Bii** | starved MBFCs | translatome | Metabolic | 55344 | None in fed, none in starved |
| **Vps25** | fed MBFCs | translatome | Metabolic | 26286 | Strong PNCN |
| **Vps25** | fed MBFCs | translatome | Metabolic | 54831 | None |
| **Not1** | fed SFCs | scRNA & translatome | Metabolic | 32836 | Moderate mid-stage death, stunted dorsal appendages |
| **PGRP-LC** | fed SFCs | translatome | NF-KB/Imd pathway | 33383 | Moderate PNCN, constricted or fused egg chambers, distended oocyte, premature dorsal appendage formation, dumpless |
| **SPE** | fed SFCs | translatome | NF-KB/Toll pathway | 33926 | Severe mid-stage death in fed, severe morphological defects |
| **PGRP-SD** | fed SFCs | translatome | NF-KB/Toll pathway | 60904 | Moderate PNCN, dumpless |
| **numb** | fed SFCs | translatome | notch pathway | 35045 | several midstage dying, distended or fused egg chambers, dumpless, stunted dorsal appendages, excessive stage 14 egg chambers (egg retention) |
| **wgn** | fed SFCs | translatome | TNF-a pathway | 55275 | None [58] |
| **Dad** | fed SFCs | scRNA & translatome | TGF-b pathway | 33759 | Dumpless, midstage death, fused egg chambers and distended SFC membranes |
| **Act79B** | fed SFCs | translatome | cytoskeletal | 36857 | None |
| **Unc-115b** | fed SFCs | translatome | cytoskeletal | 40839 | Moderate PNCN |
| **cue** | fed SFCs | translatome | cytoskeletal | 36875 | Strong PNCN |
| **Scp1** | fed SFCs | translatome | cytoskeletal | 31957 | Moderate PNCN |
| **tpnC41C** | fed SFCs | scRNA & translatome | cytoskeletal | 80889 | Moderate PNCN |
| **Tm1** | fed SFCs | scRNA & translatome | cytoskeletal | 43542 | Dumpless |
| **Hsp23** | fed SFCs & starved MBFCs | scRNA & translatome | Chaperone and heat shock proteins | 44029 | None in fed, none in starved |
| **Hsp23** | fed SFCs & starved MBFCs | scRNA & translatome | Chaperone and heat shock proteins | 82961 | None in fed, none in starved |
| **Hsp23** | fed SFCs & starved MBFCs | scRNA & translatome | Chaperone and heat shock proteins | 30541* | Moderate PNCN in fed, none in starved |
| **Npc2c** | starved MBFCs | translatome | transport/binding | 61315 | None in fed, none in starved |
| **His2B** | starved MBFCs | translatome | DNA binding | 41881 | Higher mid-stage death in fed and starved |
| **CG6403** | fed MBFCs | translatome | Unknown | 55983 | None |

Similarly, we investigated a number of genes differentially regulated in starved MBFCs. We found that ovaries from mutants for DNA binding gene *His2B,* heatshock protein *Hsp23*, and predicted triglyceride lipase *CG10163* had higher mid-stage death per ovariole than the starved matched control, while other knockdown lines had no significant detectable phenotype. (Fig 7A-7G, Table 1). Because *CG10163* is highly homologous to human lipases which have been associated with metabolic disorders such as type 2 diabetes, familial hyperlipidemia, and obesity, this candidate presents a unique opportunity to investigate crosstalk between metabolic, endocrine, and reproductive systems. In addition to observing egg chamber health in starved mutants, we investigated whether the knockdown of the candidate genes was sufficient to induce egg chamber degeneration even in well-fed flies. We found that loss of *His2B* resulted in increased mid-oogenesis death per ovariole even in well-fed flies (Fig 7G).

**Fig 7.**
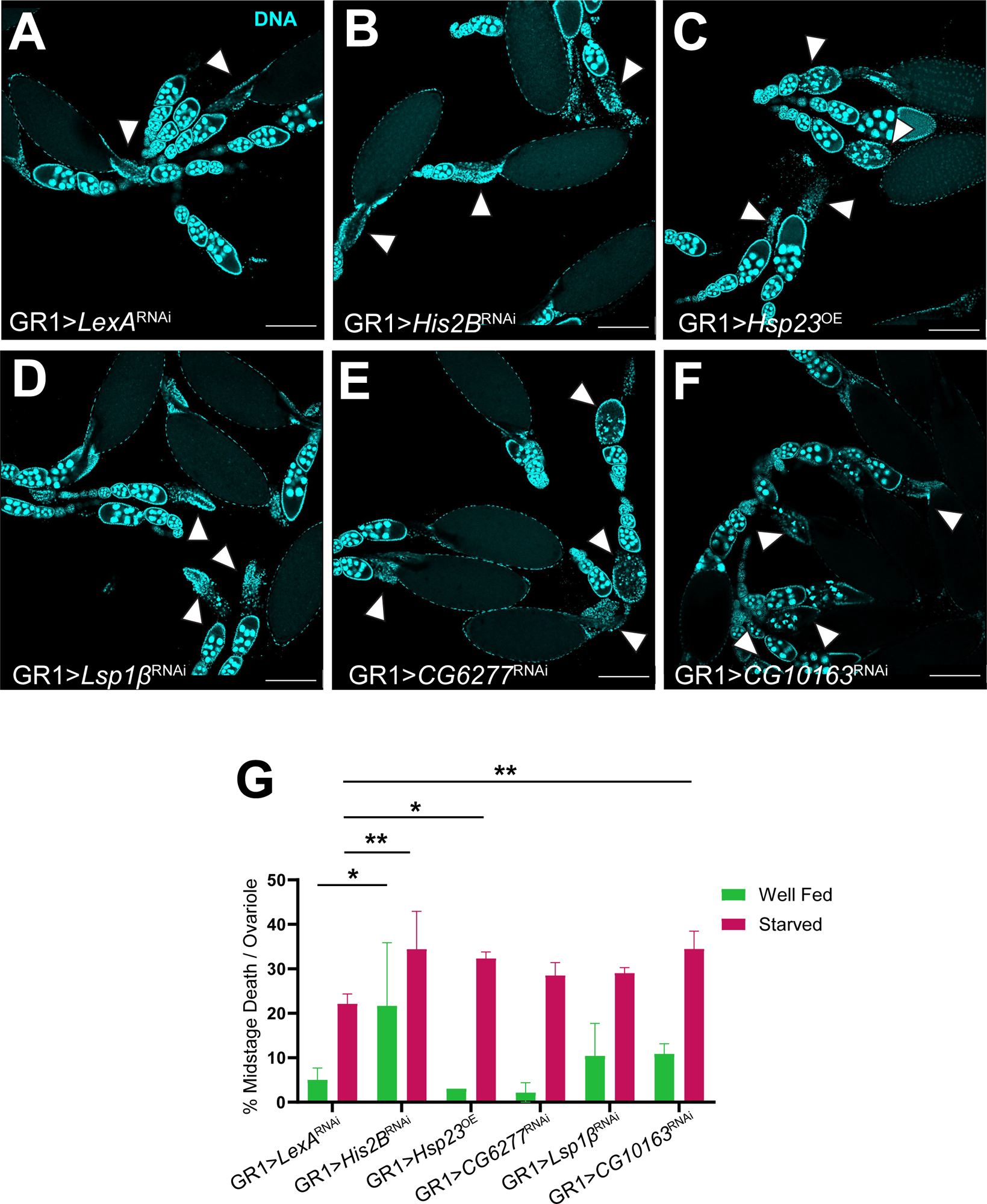
*in vivo* RNAi screening under protein-starved condition. (A-F) Egg chambers stained with DAPI (cyan) from the indicated genotypes under protein starvation. (A) *GR1>LexA RNAi* starved control shows sporadically degenerating egg chambers. (B-F) RNAi knockdowns of genes enriched in starved MBFCs. (G) Quantitative analysis of midstage degenerating egg chambers with one-way ANOVA.

## Discussion

The role of epithelial cells as non-professional phagocytes is well established. Previous studies have focused on singular models of apoptotic cell clearance. To understand how the same cellular populations can participate in two distinct death paradigms, one apoptotic, and the other non-apoptotic, we characterized the multiple levels of gene regulation required to promote these events during *Drosophila* oogenesis. Despite the shared genetic lineage and spatiotemporal proximity of well-fed MBFCs, starved MBFCs, and SFCs, we show that their translatomes and secretomes are sufficient to delineate them into distinct cellular species. We hypothesized that instantaneous cellular responses in clearance, specifically during physiological insults such as nutrient-deprivation, would be orchestrated by translational bursts, rather than by energetically expensive transcriptional responses [59]. With the availability of single-cell gene expression profiles of MBFCs and SFCs [24,25,40,60], we identified the sets of genes that are post-transcriptionally controlled. Furthermore, whole-cell RNA sequencing data from phagocytic cells often contain contaminating mRNAs from the payload of engulfed cell corpses. By combining TRAP-seq and LC-MS/MS with the GAL4-UAS system, we afford finer control over both the cell types, as well as the mRNA species or peptides being profiled. By sequencing only those mRNAs that are bound to ribosomes, we could also disregard mRNAs that are primed for degradation and discover genes that are more likely to play a causal role in promoting clearance.

With this comprehensive resource, we first recapitulated that the egg chamber matrisome is disrupted in both SFCs and starved MBFCs. This is accomplished by downregulating several components of the vitelline membrane, the chorion, and the yolk, which was captured in all three datasets. We also showed that glucose and lipid homeostasis is disrupted in the ovaries in response to nutrient deprivation, evidenced by the translational upregulation of *pepck1* and *FASN2* by starved MBFCs. These results also reassured us that we were capturing the right cell populations. Several genes with functions in cytoskeletal remodeling were identified in both death modalities, however only *Act88F* was common to them. A subset of the genes identified exclusively in the SFC translatome are known or predicted to encode proteins that bind calcium to promote myofibril assembly. *In vivo* validation of some of these candidates by RNAi revealed a strong PNCN phenotype, which supports the hypothesis that nonprofessional phagocytes, like professional phagocytes, utilize calcium signaling during cell corpse clearance [61]. We also showed for the first time that different AMPs are preferentially induced during phagoptosis and starvation-death of NCs, *Drs* in the former and *AttD* in the latter. Moreover, SFCs upregulated the translation of multiple upstream regulators of the canonical Toll signaling pathway, including PGRP-LC, PGRP-SD, and SPE. However, it remains to be known if the Toll pathway has an immunogenic role in phagoptosis. Overall, we validated 64 transgenic lines *in vivo (*Table1). Interestingly, several RNAi candidates with the GR1 driver affected NC survival rather than MBFCs. Because GR1 is not known to be active in NCs, it suggests that NC survival is determined by the health of the surrounding follicle cells.

The results presented herein also raise multiple questions. For example, we discovered that *AttD* translation is upregulated by starved MBFCs and that loss of *AttD* exacerbates egg chamber degeneration. But what role does *AttD* play during starvation-stress and what factors are required upstream to activate *AttD*? Neither the translatome nor the secretome revealed a role for canonical members of the Imd cascade in the activation of *AttD* in starved MBFCs. It would be worthwhile to test *in vivo* whether Imd pathway mutants, specifically in the transcription factor *Relish*, affect *AttD* induction during starvation-stress. Because the *AttD* genetic and amino acid sequences are a phylogenetic outgroup among *Drosophila* Attacins, it is possible that *AttD* induction has a non-immunogenic role or even results from a heretofore unknown mechanism [62]. Hedengren et al. also noted that *AttD* lacks signal peptides and hypothesize that it could have a role in intracellular defense response to pathogenic challenge [59]. Moreover, there is evidence that suggests that the vast repertoire of *Drosophila* AMPs are used combinatorially, and that their specificity is determined by the source of the infection [63]. Taken together, we hypothesize that pathogenic and non-pathogenic insults may elicit diverse AMP responses. Characterizing the substrates required for preferential AMP induction in oogenesis would have implications beyond understanding how discriminatory immune responses are regulated. Nutrient availability is a key evolutionary selective pressure that influences life-history trade-offs between survival and reproduction at the individual level. Resource allocation constraints affect not only maternal health and fecundity, but also determine the offspring size and fitness. Ultimately, the molecular differences and similarities between the two death paradigms in oogenesis highlighted in this report takes us a step closer to understanding and appreciating how cell death shapes and sustains life.

While this study is a valuable resource to generate several new hypotheses, our findings are constrained by the limited availability of samples. SFCs make up a small fraction of the egg chamber cellular demographic. In addition to low abundance of cells, capturing specific subsets of mRNAs and proteins might underrepresent species that are lowly abundant or those that occupy spatiotemporal niches, as well as those that undergo alternative modes of post-transcriptional regulation. For example, *drpr,* which has been established to be essential for engulfment and has been shown to increase in engulfing FCs by immunostaining, was not reported to be upregulated by SFCs or starved MBFCs in any of the datasets. Therefore, exploration of other means of regulation that explain the mechanism by which these notable genes are involved is warranted. Furthermore, our *in vivo* validation of the findings is constrained by the limited availability and potential imprecise targeting effects of RNAi constructs. While we expected our *in vivo* RNAi candidates to yield severe clearance defects leading to strong persistence of NCs, we only observed weak to moderate clearance defects in most candidates, in addition to a plethora of morphological defects (Table1). This could also be attributed to epistatic effects or could further indicate the presence of redundancy in clearance pathways. Additional confounding may result from the non-specificity of GR1 and PG150, as they are known to be weakly expressed in the gut and Malpighian tubules respectively and a known issue with the GAL4 system. Further experiments beyond RNAi and overexpression are required to disentangle the specific contributions of these genes and their complex interactions to promote NC death and clearance.

## Acknowledgements

We thank Drs. Samantha You, Alder Yu, and Robert Jackson, for advice on TRAP-seq and Ryan Hekman for support with tandem mass-spectrometry. Our sincere thanks to Dr. Todd Blute for microscopy support and Cameron Dixon and Yuanhang Zhang for technical assistance. We thank the Bloomington *Drosophila* Stock Center, the *Drosophila* Genome Resource Center, and Developmental Studies Hybridoma Bank for strains and reagents. We also thank Dr. Norbert Perrimon, and other current and previous members of the McCall lab for helpful discussions.

## Materials and Methods

### Fly husbandry

All flies were obtained from Bloomington Stock Center, or other laboratories in the Drosophila community (Table 1). *GR1-GAL4* (gift from Dr. Trudi Schüpbach) was utilized to drive expression in follicle cells, *PG150-GAL4* (gift from Dr. Ellen LeMosy) was used to driveexpression in stretch follicle cells [64,65]. *GR1* is expressed in all FCs beginning in stage 3 egg chambers [6]. *PG150* is expressed in the SFCs and centripetal cells. *UAS-Luciferase RNAi* (Bloomington stock #31603) or *UAS-LexA RNAi* (Bloomington stock #67946) was used as a control for all RNAi experiments [66]. All flies were raised on standard molasses/yeast/agar/cornmeal food. Long term fly stocks were stored at 18 °C. Active stocks were raised at 25 °C. Crosses were performed at 25 C°.

## Translatome library preparation and bulk RNA-Seq

### Fly abdomen collection

To collect enough ovaries for RNA sequencing and prevent RNA degradation, whole fly thorax and abdomens were collected from samples containing approximately 100 flies. Flies of the appropriate genotype were transferred to a 15 mL conical tube and submerged in liquid N2 for 1 min. The conical tube was then vortexed vigorously for 20 s to break flies apart and tissues were collected over a set of sifters to obtain thoraxes and abdomens. Once samples were processed, they were transferred to a 1.5 mL microcentrifuge tube and stored in a -80°C freezer for up to one month.

### Immunoprecipitation of ribosome-bound mRNAs

Ribosome immunoprecipitation was performed as per [67]. Fresh magnetic beads were prepared from the Dynabeads antibody coupling kit (Life Technologies) and conjugated to the19C8 anti-EGFP antibody (Memorial Sloan Kettering). Abdomens of 100 flies were homogenized on ice in 400 μl of homogenization buffer (9.1 mL nuclease free H2O (Ambion), 200 *µ*L 1M HEPES-KOH pH 7.4 (USB), 750 *µ*L 2M KCl (Ambion), 50 *µ*L 1M MgCl2 (Ambion), 1 Complete Mini EDTA-Free Protease Inhibitor Cocktail Tablet (Roche), 5 *µ*L 1M DTT (Sigma), 10 *µ*L Super RNasin (Ambion), and 10 *µ*L of freshly prepared 1000 mg/mL cycloheximide (Sigma)). Homogenized fly samples were then centrifuged for 30 min at 20,000g at 4 °C. The supernatant was transferred to new pre-chilled 1.5 mL tubes. A 1:1 mixture of 300 mM 1,2-diheptanoyl-sn-glycero-3-phosphocholine (DHPC, Avanti) and 10% Igepal (USB) was added to the to the supernatant in a ratio of 1 part mixture: 4 parts supernatant. The solutions were mixed by inversion, incubated on ice for 5 min, then centrifuged for 5 min at >20,000g at 4 °C. Cleared lysate was added to an aliquot of the bead suspension and incubated for 1 hr at 4 °C with end over end rotation. The mixture was briefly centrifuged then magnetized for 2 min. After removing the buffer, beads were rinsed 6 times in 1 mL 0.35M KCl Wash Buffer (7.1 mL nuclease-free H2O (Ambion), 200 *µ*L 1M HEPES-KOH pH 7.4 (USB), 1.75 mL 2M KCl (Ambion), 50 *µ*L 1M MgCl2 (Ambion), 1 mL 10% Igepal (USB), 5 *µ*L 1M DTT (Sigma), and 50 *µ*L of freshly prepared 1000 mg/mL cycloheximide (Sigma)). After final rinse, beads were resuspended in 200 μl of nuclease-free water (Ambion) and RNA was extracted with TRIzol (Invitrogen). RNA quality was assessed with an Agilent RNA 6000 Bioanalyzer.

### cDNA library production and sequencing

The Illumina TruSeq RNA Sample Preparation v2 was used to convert 0.1 – 4 *µ*g of precipitated mRNA to cDNA in preparation for RNAseq. Poly-T oligo-attached magnetic beads were used to isolate mRNA and the mRNA was fragmented and primed with random hexamers for cDNA synthesis using SuperScript II. Second strand cDNA synthesis was performed and in preparation for attaching the Illumina adaptor sequences, a single A nucleotide was added to the 3’ ends of the blunt fragments. Samples were run on an Agilent BioAnalyzer (BU Microarray core). Once the samples were validated by the core, they normalized, pooled, and sequenced on an Illumina NextSeq500. The sequencer was instructed to run 400 million 75bp single end reads.

### TRAP-seq QC, differential expression analysis, and functional enrichment

RNA-seq data from biological triplicates corresponding to well-fed and starved MBFCs (*GR1-GAL4 > UAS-RpL10a-GFP*), and well-fed SFCs (*PG150-GAL4 > UAS-RpL10a-GFP*) were assessed for sequence quality using FastQC v0.11.7. Illumina sequencing adapters were computationally removed by the Boston University microarray sequencing core facility, where the sequencing was performed. Samples were evaluated for per-base Phred score and for overrepresented sequences. Reads that passed QC evaluation were aligned to the *Drosophila melanogaster* reference genome BDGP6.32 release 104 using Salmon v1.1.0 in the quasi-mapping mode. This step was parameterized to automatically infer sequence library type using the -l A flag, correct for fragment-level gc-bias, as well as perform selective alignment using – validatemappings.

The aligned reads in the salmon quant file format were then imported into DESeq2 to identify differentially translated genes. Genotype (MBFC vs. SFC) and treatment (well-fed vs. starved) were both used as factors in the DESeq2 linear model design [68]. Log fold change estimates were obtained after applying the ApeGLM method for effect size shrinkage [69]. ClusterProfiler v4.2.2 and PANGEA were used to identify the pathways and gene ontology terms enriched in both comparisons [70,71]. GLAD webtool was used to obtain additional functional annotations [72].

## Secretome Sample Preparation and Liquid Chromatography-Tandem Mass Spectrometry

### Molecular cloning of *UAS-ss-HRP-KDEL-V5*

pDisplay-ss-V5-HRPKDEL was obtained from Dr. Alice Ting [30].The insert was PCR-amplified and cloned into the pENTR/D-TOPO cloning kit (Invitrogen) and pTW vector (*Drosophila* Genome Resource Center, RRID:DGRC_1129)) using Gateway cloning (Invitrogen). The plasmid was purified by Qiagen Midiprep kit, confirmed by sequencing and sent to BestGene (Chino Hills, CA) for injection into *Drosophila* embryos.

### Protein biotinylation using HRP-KDEL

Freshly dissected ovaries were incubated in 300 μL of 500 uM biotin phenol for 30 minutes at room temperature rotating. Samples were then rinsed with 1X PBS twice and the biotinylation reaction was initiated by adding 1 mM H_2_O_2_ in PBS to the samples for 1 minute and rotating at room temperature. Ovaries were quickly washed with quencher solution (10 mM sodium ascorbate, 5 mM Trolox (Sigma-Aldrich), 10 mM sodium azide, then lysed in 100 μL RIPA buffer with quencher solution for 5 min on ice. RIPA buffer was composed of: 50 μL 1M Tris-HCl, 150 μL 5M NaCl, 50 μL of 10% SDS, 250 μL of 10% Sodium Deoxycholate, 500 μL of 10% TritonX-100, 50 μL of 100X Protease Inhibitor (Sigma-Aldrich – P8849), 50 μL of 100 mM PMSF, 3.550 mL of diH20. Tissue was homogenized by motorized pestle and centrifuged at 16.1g for 10 min at 4°C. Clarified sample (clear middle layer) was transferred to a new tube and snap frozen in liquid nitrogen. This protocol was adapted from [73,74].

### Biotinylated protein pull-down

Frozen protein samples were thawed on ice. Meanwhile, 50 μL streptavidin magnetic beads (Pierce 88817) were washed with 1 mL of RIPA lysis buffer twice. The beads were subsequently incubated with 90 μL of protein lysate (about 550 μg of protein) and an additional 500 μL of RIPA buffer was added to facilitate rotation for 1 hour at room temperature. Beads were pelleted on a magnetic rack and the supernatant (flow-through) was collected on ice. After each wash, the magnetic beads were transferred into new tubes and washed twice with 1 mL RIPA buffer, once with 1 mL KCl, once with 1 mL 0.1 M Na_2_CO_3_, once with 1 mL 2 M Urea in 10 mM Tris-HCl (pH 8), and twice with 1 mL RIPA buffer. For the proteomic analysis, RIPA buffer was removed, and the beads snap frozen in liquid nitrogen and stored at -80°C. Otherwise, protein was eluted by boiling each sample in 30 μL of 3X protein loading buffer (containing DTT, Pierce) supplemented with 2 mM biotin (Sigma-Aldrich) for 10 min. After boiling, the beads were pelleted with the magnetic rack and the eluate collected for analysis by Western blot. Biotinylated proteins were probed on western blot with streptavidin-HRP (Thermo Fisher Scientific).

### Western blotting

Protein was isolated by lysing whole ovaries in 100 μL fresh ice-cold RIPA buffer for 5 minutes. After the 5-minute incubation, tissue was disrupted by using a motorized pestle until clumps were broken down (about 30 seconds). Samples were centrifuged for 10 minutes at 4°C at 16.1 relative centrifugal force (RCF). The clarified sample (clear middle layer only) was transferred to a new tube. Protein that was to be analyzed later was snap frozen in liquid nitrogen and stored at -80°C. Laemmli loading buffer with 2-mercaptoethanol (BME, Bio-Rad) was added to the protein samples (1:3), boiled at 95°C and then immediately stored on ice. Samples were run on a 10% resolving gel, transferred to nitrocellulose and protein was detected by enhanced chemiluminescence (ECL, Thermo Fisher Scientific).

### On-bead trypsin digestion and LC-MS/MS

Beads from biotinylated protein pull-down were washed with 100 mM triethylammonium bicarbonate. Peptides were eluted from beads by on-bead trypsin digestion with 1μg Trypsin (Pierce) in 100 mM triethylammonium bicarbonate overnight rotating at 37 °C. Peptides were desalted using C18 ZipTip (Millipore) and subjected liquid chromatography coupled to tandem mass spectrometry on a Q Exactive HF-X (Thermo Fisher Scientific). Data-dependent fragmentation used collision-induced dissociation. RAW files were searched using MaxQuant under standard settings using the UniProt *Drosophila melanogaster* database, allowing for two missed trypsin cleavage sites, variable modifications for N-terminal acetylation, and methionine oxidation. Candidate peptides and protein identifications were filtered on the basis of a 1% false discovery rate.

### Secretome QC, differential abundance estimation, and functional enrichment

Raw intensities from the MaxQuant spectral database search engine were imported into R for downstream analysis. QC metrics such as number of unique peptides identified, percent of contaminants and reverse decoys detected, and total sum of intensities at the replicate level and condition level were computed and visualized using artMS v1.12.0. Spurious hits such as reverse decoys and potential contaminants were removed. R package DEP v1.16.0 was used to remove proteins not identified in both replicates simultaneously and variance stabilizing transformation was applied to normalize the intensities. Missing values were imputed using the k-nearest neighbors imputation method, followed by differential abundance estimation and multiple hypothesis testing correction in limma, all of which were implemented in DEP.

## Single cell RNA-seq data integration

### Integration of Rust et. al. 2020 and Jevitt et. al. 2020

Previously published single cell RNA-sequencing datasets of the *Drosophila melanogaster* ovary from Rust et. al. and Jevitt et. al. aligned using CellRanger were obtained from the Gene Expression Omnibus database (GEO: GSE136162 and GSE146040 respectively). Seurat v4.1.1 was used to perform quality control, pre-processing, data integration, and differential expression analysis. Low quality cells were removed from individual datasets as described in the original publications prior to integration. The datasets were then individually normalized using sctransform implemented in Seurat using the glmGamPoi estimator for features that are present in at least 1 cell. Seurat RPCA (Reciprocal PCA) was used to integrate the two normalized datasets using an augmented feature list containing 3000 top ranking features from *SelectIntegrationFeatures* (Seurat) and a list of known MBFC and SFC marker genes. 100 neighbors (*k.anchors*) were used to achieve sufficient strength of integration in the *FindIntegrationAnchors* (Seurat) step [31]. Principal components were estimated for the integrated dataset and the top 30 dimensions were used to obtain the UMAP embeddings. New clusters were identified using the Louvain algorithm implemented in Seurat with the resolution set to 5. Cell identity labels from the original publications were retained in the integrated dataset to verify that clusters in the integrated dataset contain cells from analogous tissues in the original datasets.

### Single Cell RNA-seq SFC and MBFC module score estimation, reclustering, and differential expression analysis

Seurat *AddModuleScore* was used to identify MBFC and SFC populations equivalent to those in the translatome and secretome datasets. To identify SFCs, previously published SFC-enriched genes *cv-2, dpp, eya, Past1, peb, drpr, puc, trol, kay*, *Vha16-1, Vha100-2* were used to compute an “SFCScore”. Similarly, to identify MBFCs, we scored cells using canonical markers such as *Yp1*, *Sox14*, and *br*. Cells with SFCScore >1 were designated as SFCs and cells with MBFCScore > 2 were designated as MBFCs. The integrated dataset was then subset to MBFC and SFC cells and reclustered. Differentially expressed genes between MBFC and SFC cells were identified using Seurat functions *PrepSCTFindMarkers* and *FindMarkers*, using the DESeq2 test with logfc.threshold parameter set to 0.25.

## DAPI and antibody staining

For the well-fed experimental condition, flies were aged 3 days at 25 °C on standard cornmeal, molasses, and agar mixture, and then fed fresh yeast paste every 24 hours for 2 consecutive days before abdomens were dissected. For the protein starvation condition, flies were transferred for an additional day to apple juice agar lacking yeast. Ovaries were harvested in fresh 1X PBS and fixed at room temperature for 20 minutes in a solution of 4% paraformaldehyde (less than a week old), and 1X PBS, followed by two successive rinses in 1X PBT (1X PBS with 0.1% Triton X-100). The tissues were then washed 3 times in 1X PBT, each wash lasting 20 minutes. Following a rinse with 1X PBS, the tissues were incubated overnight at 4 °C in 2 drops of Vectashield + 4’,6-diamidino-2-phenylindole dihydrochloride (DAPI) (Vector Laboratories) prior to mounting on slides.

Antibody staining was performed on ovaries following fixation and washing in PBT, as described above. Tissues were blocked in PBTG (PBT containing 1.5% normal goat serum) for 1 hour and then incubated in Discs large antibody (43F, DHSB), diluted 1/500 in PBTG, overnight at 4 °C with agitation. The tissues were then washed in 4 times in PBT over 2 hours. The secondary antibody was Goat anti-mouse-Cy3 (Jackson Labs) diluted in PBTG (1:200) and incubated for 2 hours at room temperature. The tissues were then washed 4 times in PBT and mounted in DAPI (Vector Labs).

### Imaging, quantification of mutant phenotypes, and statistics

DAPI stained tissues were imaged on the Olympus BX60 upright fluorescence microscope or on the Olympus FV10i confocal microscope or the Nikon C2si confocal microscope. Image stacks were processed in FIJI and Adobe Illustrator. Stage 14 egg chambers were identified by the presence of a pair of fully formed dorsal appendages. The effect of disruption of phagoptosis of NCs resulting from RNAi perturbation of specific genes was quantified by the persistence of NC nuclei in stage 14 egg chambers. Persisting NC nuclei (PNCN) were visually identified and scored by binning the number of PNCN in each stage 14 egg chamber into 6 classes of phenotype severity, starting from egg chambers with 0 PNCN, and subsequently 1-3, 4-6, 7-9, 10-12, 13-15 PNCN. Egg chambers undergoing midstage death were identified by NC nuclear condensation and fragmentation as well as FC membrane enlargement (utilizing α-DLG staining). Midstage dying egg chambers were visually identified and quantified along with the number of germaria present per slide. The percentage of midstage death per ovariole was calculated by dividing the number of midstage dying eggs by the number of germaria and multiplied by 100. Two-way ANOVA was used to calculate the statistical difference between both starved samples to the starved control and fed samples to the fed control. Those that had a p-value of < 0.05 were considered statistically significant.

## Code and Data Availability

All scripts used for data quality control, analysis, and visualization can be accessed via GitHub https://github.com/McCallLabBU/multimodal_nurse_cell_clearance

---

**S1 Fig.**
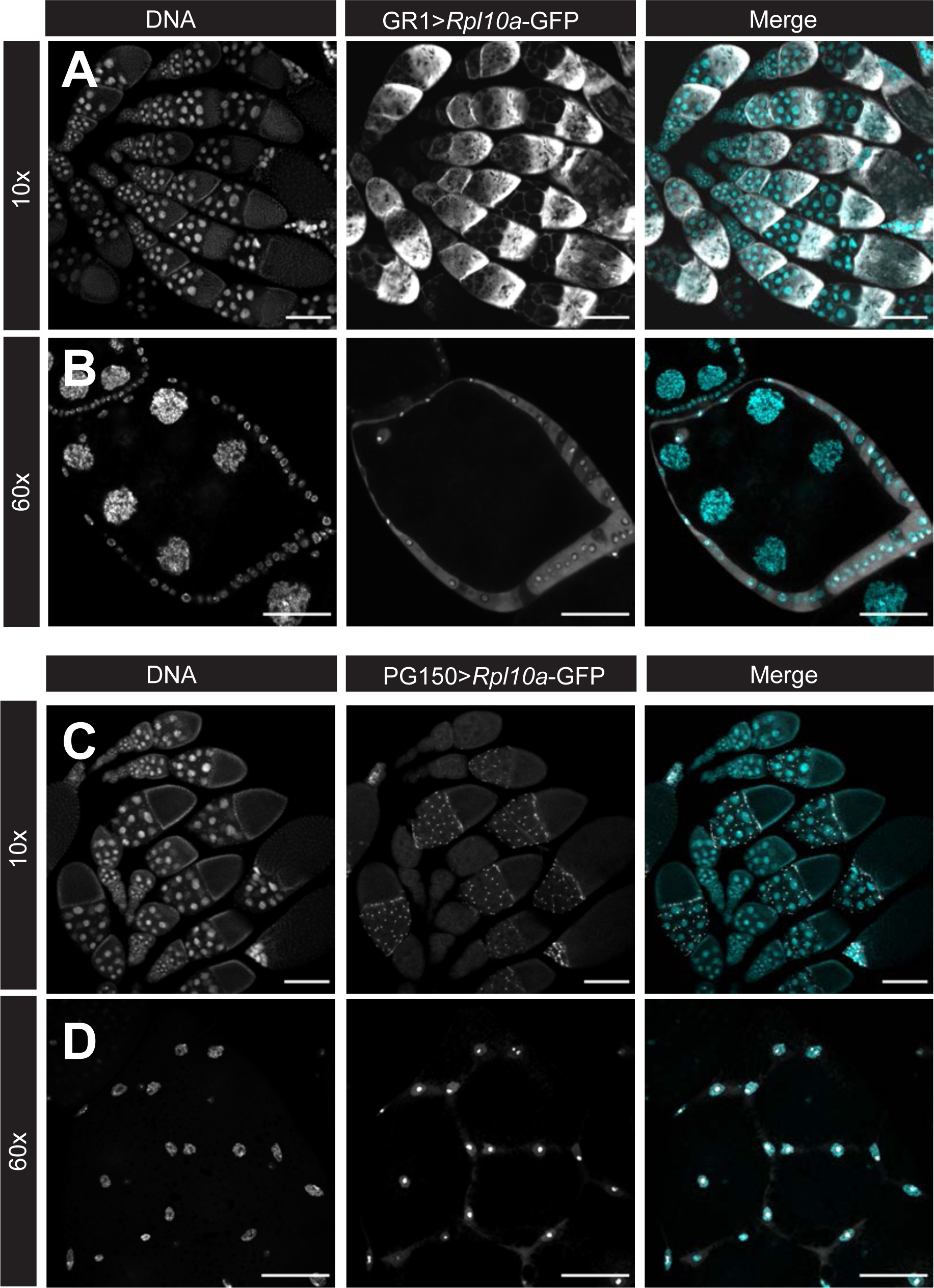
Characterization of the Rpl10a-GFP construct for TRAP-seq. (A-D) Egg chambers of indicated genotypes stained with DAPI. (A) Expression of *GR1 > Rpl10A-GFP* in MBFCs. Scale bars= 200µ (B) Expression of *GR1 > Rpl10A-GFP* in MBFCs. Scale bars= 50µ (C) Expression of *PG150 > Rpl10A-GFP* in SFCs. Scale bars= 200µ (D) Expression of *PG150 > Rpl10A-GFP* in SFCs. Scale bars= 50µ.

**S2 Fig.**
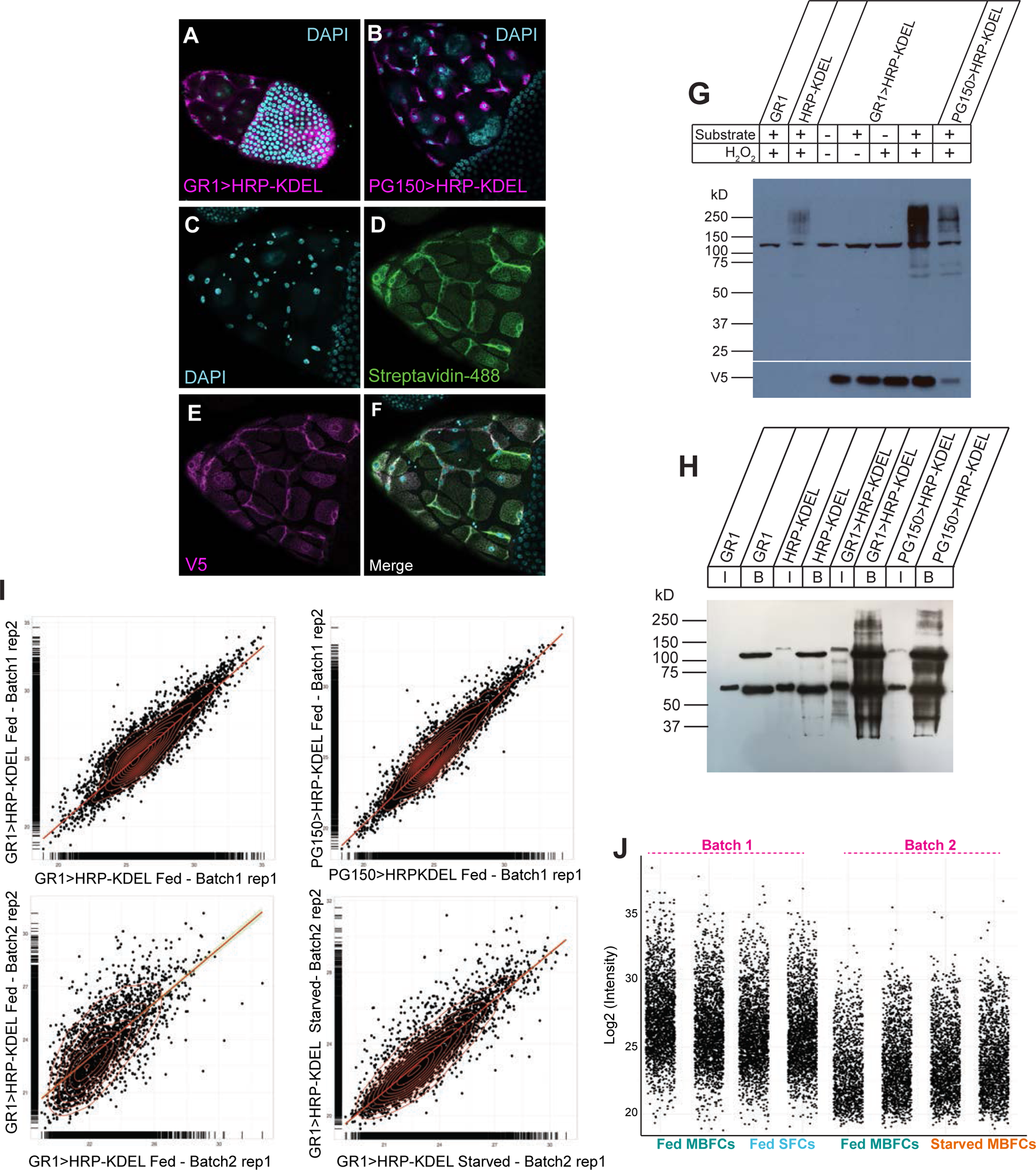
Characterization of the HRP-KDEL construct for LC-MS/MS. (A-B) Egg chambers of indicated genotypes stained for DAPI (cyan) and anti-V5 (magenta) to detect HRP-KDEL. (A) *GR1 > UAS-HRP-*KDEL-V5 stage 8 egg chamber shows HRP-KDEL in all follicle cells. (B) *PG150 > UAS-HRP-KDEL-V5* stage 10 egg chamber shows HRP-KDEL in stretch follicle cells only. (C-F) *PG150 > UAS-HRP-KDEL-V5* stage 11 egg chamber with (C) DAPI, (D) biotinylated proteins (streptavidin-488, green), and (E) HRP-KDEL (anti-V5, magenta). (F) Protein biotinylation pattern is restricted to cells expressing HRP-KDEL. (G) (Top) Western blot analysis of lysate (30 μg per lane) from ovaries of indicated genotypes (GR1 – *GR1-GAL4*, HRP-KDEL – *UAS-HRP-KDEL-V5*, GR1>HRP-KDEL – *GR1-GAL4; UAS-HRP-KDEL-V5*, and PG150>HRP-KDEL – *PG150-GAL4; UAS-HRP-KDEL-V5*) incubated with or without substrate (biotin-phenol) and H_2_O_2_. *GR1>HRP-KDEL* and *PG150>HRP-KDEL* sample with both substrate and H_2_O_2_ have many biotinylated proteins as detected by streptavidin-HRP. (Bottom) α-V5 staining confirms HRP-KDEL expression in ovary tissue. (H) Western blot probed with streptavidin-HRP. Biotinylated proteins before (I-input, 30 μg) and after (B-beads, all of eluted protein) streptavidin enrichment. Genotypes as in G. (I) Scatterplot of peptide log2 intensity values of one replicate against another in each condition. (J) Distribution of log2 peptide intensity values in each replicate for all conditions.

**S3 Fig.**
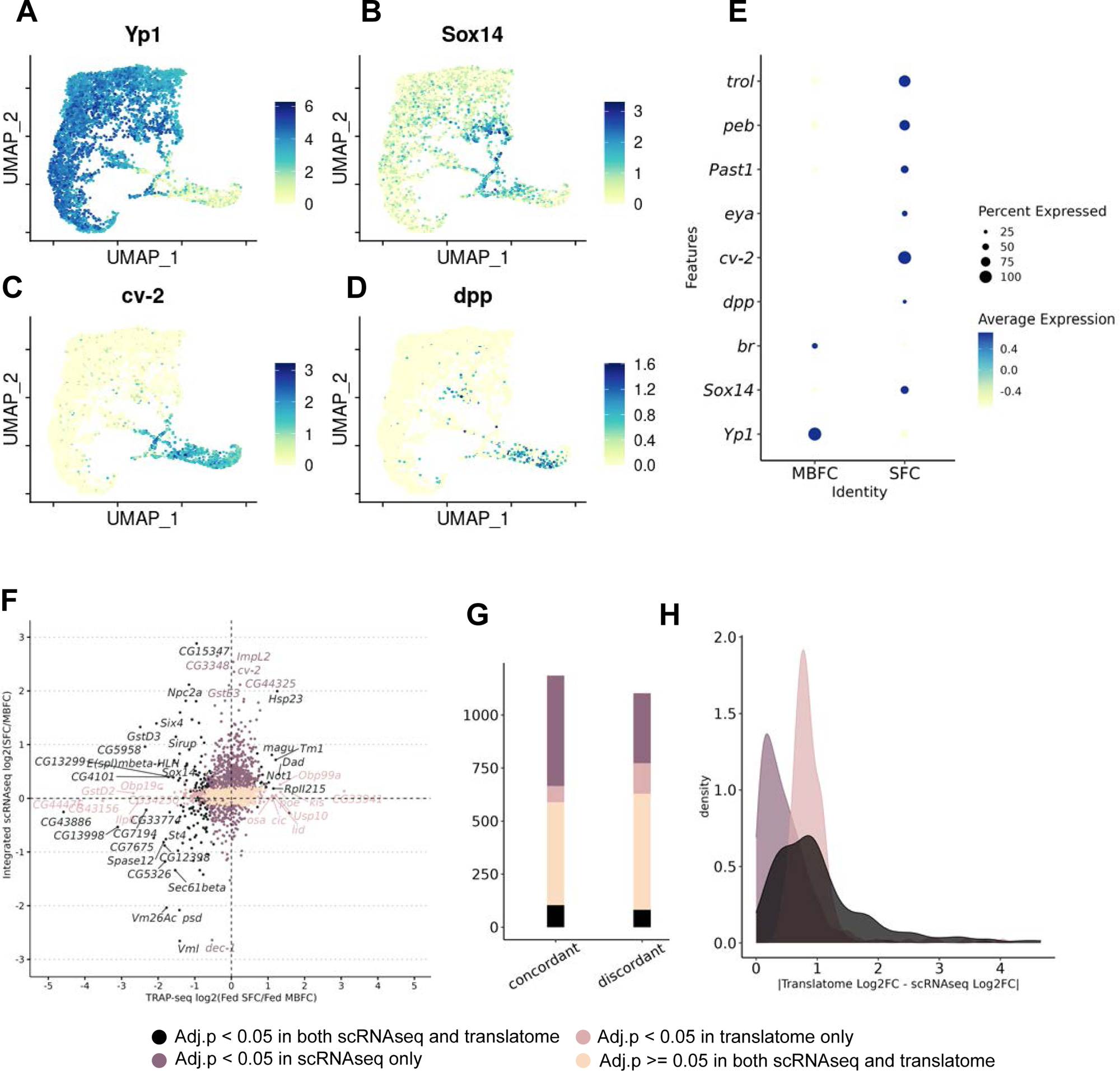
Integration of scRNAseq and comparison to the translatome. (A-D) Expression levels of canonical MBFC and SFC markers (E) Dot plot showing average expression and the percent of cells expressing each canonical marker in SFCs and MBFCs (F) Scatterplot of Log2 Fold Change values of differentially translated genes in the translatome (x-axis) and Log2 Fold Change values of differentially expressed genes in the integrated single cell RNAseq MBFC-SFC subset (y-axis). Positive values on both axes indicate upregulation in SFCs, when compared to the MBFC baseline (denominator, negative values on both axes). The points are colored by whether the L2FC differential is significant at the adjusted p-value threshold of 0.05 in both, either, or neither comparison. (G) Summary of total number of genes in each category grouped by the direction of L2FC differential in scRNA-seq and translatome datasets. (H) Distribution of absolute value of differences in L2FC in translatome and scRNAseq.

## Supplemental Tables

**S1 Table. GO-terms enriched in phagoptosis and starvation-death identified in the translatome.**

Summary of the number of genes congruently (up or down) and uniquely regulated between SFCs and starved MBFCs and the GO terms enriched in each category, along with the adjusted p-value.

**S2 Table. Differentially regulated candidates (Log fold change and FDR adjusted p-values) in phagoptosis across the translatome, secretome, and the integrated scRNA-seq datasets.**

Log fold change and FDR adjusted p-values obtained from DESeq2 (translatome, scRNA-seq) or DEP (secretome) are summarized for the SFC vs. fed MBFC comparison.

**S3 Table. Differentially regulated candidates in starvation-death across the translatome, secretome, and the integrated scRNA-seq datasets.**

Log fold change and FDR adjusted p-values obtained from DESeq2 (translatome, scRNA-seq) or DEP (secretome) are summarized for the starved MBFC vs. fed MBFC comparison.

**S4 Table. Consolidated summary of candidates differentially expressed in the translatome in both phagoptosis and starvation-death.**

Log fold change and FDR adjusted p-values obtained from DESeq2 are summarized for each gene in both SFC vs fed MBFC comparison and starved MBFC vs. fed MBFC comparison. Genes are categorized as congruently up or down-regulated or as uniquely regulated in phagoptosis or starvation based on the fold change and FDR adjusted p-values across both comparisons.

